# Identification of miR-187 as a modulator of early oogenesis and female fecundity in medaka

**DOI:** 10.64898/2026.01.09.698593

**Authors:** Marlène Davilma, Stéphanie Gay, Manon Thomas, Sully Mak, Fabrice Mahé, Laurence Dubreil, Jérôme Montfort, Aurélien Brionne, Julien Bobe, Violette Thermes

**Affiliations:** INRAE, LPGP, 35000, Rennes, France; Université de Rennes, CNRS, IRMAR - UMR 6625, F-35000, Rennes, France; Oniris, INRAE, APEX, PAnTher, 44300 Nantes, France

**Keywords:** Ovary, Fish reproduction, MicroRNAs, CRISPR/Cas9, 3D imaging, Next generation sequencing

## Abstract

MicroRNAs (miRNAs) are known regulators of ovarian function in vertebrates, yet their physiological roles in fish reproduction remain poorly understood. Here, we identified miR**-**187 as one of the most ovarian-enriched miRNAs in medaka (*Oryzias latipes*) and we uncovered its function *in vivo* using CRISPR/Cas9-mediated gene inactivation. miR-187-3p is expressed in oocytes and granulosa cells in the ovary, and in discrete brain areas. Its loss-of-function leads to a significant reduction in female fecundity. High-resolution 3D imaging of whole ovaries revealed that *mir-187* mutants accumulate early stage I follicles and show reduced progression to later stages, indicating a defect in early follicle recruitment and growth. Transcriptomic profiling of mutant ovaries revealed extensive gene-expression remodeling, including downregulation of key regulators of steroidogenesis, Wnt/β-catenin signaling, and TGF-β pathways, and upregulation of genes associated with early follicle activation and immature somatic cell states. Using an expression-based target-prediction pipeline, we identified several putative miR-187-3p targets, including *nr6a1a* (*gcnf*) and *dpagt1*, two genes previously implicated in oocyte differentiation and female fertility in mammals. Together, our results demonstrate that miR-187 acts as a previously unrecognized regulator of early folliculogenesis and female reproductive capacity in medaka, expanding the repertoire of miRNAs with essential *in vivo* roles in teleost oogenesis and female fecundity.

**AUTHOR SUMMARY:** Successful fish reproduction depends on the continuous production of oocytes within the ovary. This process depends on the coordinated development of germ cells and the supporting somatic cells. Although microRNAs (miRNAs) are known to post-transcriptionally regulate gene expression, their physiological roles in the fish ovary and female reproduction remain poorly understood. In this study, we used the medaka to investigate the function of miR-187, a miRNA strongly enriched in the ovary. By generating a CRISPR/Cas9 knockout line, we showed that females lacking miR-187 produce fewer eggs and display defects in early follicle development. Using whole-ovary 3D imaging, we found that mutant females accumulate early follicles that fail to progress through normal growth stages. Transcriptomic analyses revealed broad gene-expression changes affecting pathways involved in follicle activation, somatic cell maturation, and ovarian signaling. Our results demonstrate that miR-187 is a previously unrecognized regulator of early folliculogenesis and female fecundity in medaka. This work expands the landscape of miRNA-mediated regulation in the ovary and highlights the importance of post-transcriptional mechanisms in controlling reproductive success in teleosts.

**Grant support:** The DYNAMO project (French National Research Agency, ANR-18-CE20-0004).

The OVOPAUSE project (French National Research Agency, ANR-22-CE45-00017-02).

The IMMO project (INRAE Metaprogramme DIGIT-BIO).

## INTRODUCTION

Oocyte development, and ultimately reproductive success, relies on the coordinated differentiation and activity of several cell lineages in the ovary. In teleost fish, oocyte formation and growth require extensive cellular remodeling and tightly coordinated interactions with surrounding differentiating somatic follicular cells, including granulosa and theca cells (1). These processes are controlled by complex regulatory networks integrating transcriptional, endocrine and paracrine signals (2). In addition to these well-characterized mechanisms, post-transcriptional regulation has emerged as important modulators of gene expression in the fish ovary, notably through the action of microRNAs (miRNAs), small non-coding RNAs approximately 22 nucleotides in length, that modulate mRNA stability and translation (3–5).

Despite their prominent expression in the ovary, the biological importance of individual miRNAs is particularly difficult to infer from sequence features and expression patterns alone. Genetic studies in invertebrates have shown that loss of individual miRNAs often results in viable and incompletely penetrant phenotypes at the organ or organism level under standard laboratory conditions, despite being biologically meaningful in specific contexts (6). However, many conserved miRNAs have important biological functions in vertebrates, including mammals and fish, with phenotypes that can be severe, context-dependent, or revealed only through detailed genetic or physiological analyses (7). These observations reflect the general properties of miRNAs as modulators embedded within robust gene regulatory networks, where they contribute to developmental stability, buffering of gene expression noise, and coordination of functionally related targets (8). Accordingly, miRNA-mediated repression typically induces modest changes in target gene expression (often limited to a few percent and rarely exceeding 50%), which can be attenuated by compensatory mechanisms at the cellular or tissue level. In addition, although a single miRNA can interact with hundreds or even thousands of mRNA binding sites, only a subset of these miRNA-mRNA interactions is expected to generate a readily detectable phenotype under standard laboratory conditions (9).

Thanks to next-generation sequencing approaches, the repertoire of ovarian miRNAs in teleosts is now relatively well characterized, with approximately 300 miRNA genes in teleosts (10). On the basis of this extensive catalog, numerous *in vitro* functional studies have proposed potential roles for miRNAs in ovarian physiology, including oocyte and follicle maturation, granulosa cell proliferation or survival, and steroidogenesis. However, most of these hypotheses rely on computational target predictions and *in vitro* gain- or loss-of-function assays performed in primary ovarian cell cultures (3,5,11,12). In contrast, only a limited number of miRNAs have been functionally characterized *in vivo* in fish. Among them, miR-202 stands out as a key miRNA necessary for ovarian functions and female reproduction. miR-202 is specifically enriched in fish ovary and is required for early follicular growth in medaka, as KO females exhibit severely reduced fecundity (13,14). Another notable example is the miR-200 cluster, which is highly expressed in brain and controls final oocyte maturation in zebrafish through regulation of pituitary gonadotropins. Deletion of miR-200s results in sterile females (15). Thus, although miRNAs exert modest regulatory effects, some are essential for female reproductive success, raising the question of whether additional miRNAs among the teleost repertoire play critical roles in ovarian function and induce detectable phenotype.

Here, we used the medaka fish as a model to investigate miRNA-mediated control of reproductive capacities in teleosts. We focused on miRNAs specifically enriched in the ovary, based on the hypothesis that they play a role in ovarian functions. By exploiting available small-RNA sequencing datasets, we identified *mir-187* as a strong candidate and first characterized its ovarian and brain expression patterns. We then conducted an *in vivo* functional analysis combining CRISPR/Cas9-mediated loss-of-function, 3D imaging and transcriptomic profiling to determine the role of miR-187 in oocyte development and female reproductive capacity.

## RESULTS

### Identification of ovarian-enriched miRNAs

To identify miRNAs potentially involved in intra-ovarian regulation and fish reproduction, we searched for medaka miRNAs that are specifically enriched in the ovary compared to other tissues. We analyzed available small RNA-seq datasets from a panel of 11 adult tissues (brain, eyes, fins, gills, muscle, heart, kidney, intestine, liver, testis, and ovary)(10,14). We retained 388 miRNAs with lengths ranging from 17 to 25 nucleotides and with a minimum expression level of 10 reads-per-million reads (rpm) across all tissues. This relative abundance in each tissue was used to calculate an Organ Enrichment Index (OEI, Ludwig et al., 2016) (Fig 1A). Similar to the Tissue Specificity Index (TSI) used for mRNAs (18), the OEI ranges from 0 to 1. A score of 0 denotes uniform relative expression across all tissues, characterizing ‘housekeeping’ miRNAs, whereas a score of 1 indicates a markedly higher relative expression in one or few tissues, defining ‘organ-specific enriched’ miRNAs. The OEI was calculated for each of the 388 miRNAs. Three miRNAs (0.8% of all analyzed miRNAs) exhibited similar relative expression across tissues (OEI < 0.3), while a total of 180 miRNAs (46.4% of all analyzed miRNAs) were found to be highly enriched in one or few tissues (OEI > 0.85). Relative expression profiles of the 180 enriched miRNAs could be grouped into several clusters (Fig 1B), including 2 clusters of miRNAs predominantly enriched in the ovary. Cluster 1 consisted in 8 miRNAs (miR-7552b-3p, miR-730-5p, miR-153c-3p, miR-153c-5p, miR-7552b-5p, miR-727b-5p, miR-728b-3p, and miR-728b-5p), with some of them also enriched in brain and intestine, and all located within a single genomic locus. However, their relative expression in the ovary was low, not exceeding 450 rpm (Fig 1C). Cluster 2 consisted in 10 miRNAs (miR-187-3p, miR-187-5p, miR-430d-4-3p, miR-430a-3p, miR-430c-3p, miR-430b-3p, miR-430d-3p, miR-202-5p, miR-202-3p and miR-430d-2-5p), with some of them also enriched in testis (Fig 1D). Among them, miR-202-5p exhibited the highest relative abundance (over 200,000 rpm), followed by miR-187-3p (around 1,400 rpm), both of which are located within a single genomic locus. Members of the miR-430 family displayed a much lower abundance (less than 400 rpm). While miR-202 is already well established as a key regulator of oogenesis in medaka (14), and the miR-430 family is well-established for its role in early embryonic development (19), the poorly characterized miR-187 was retained for further analysis.

**Fig 1.**
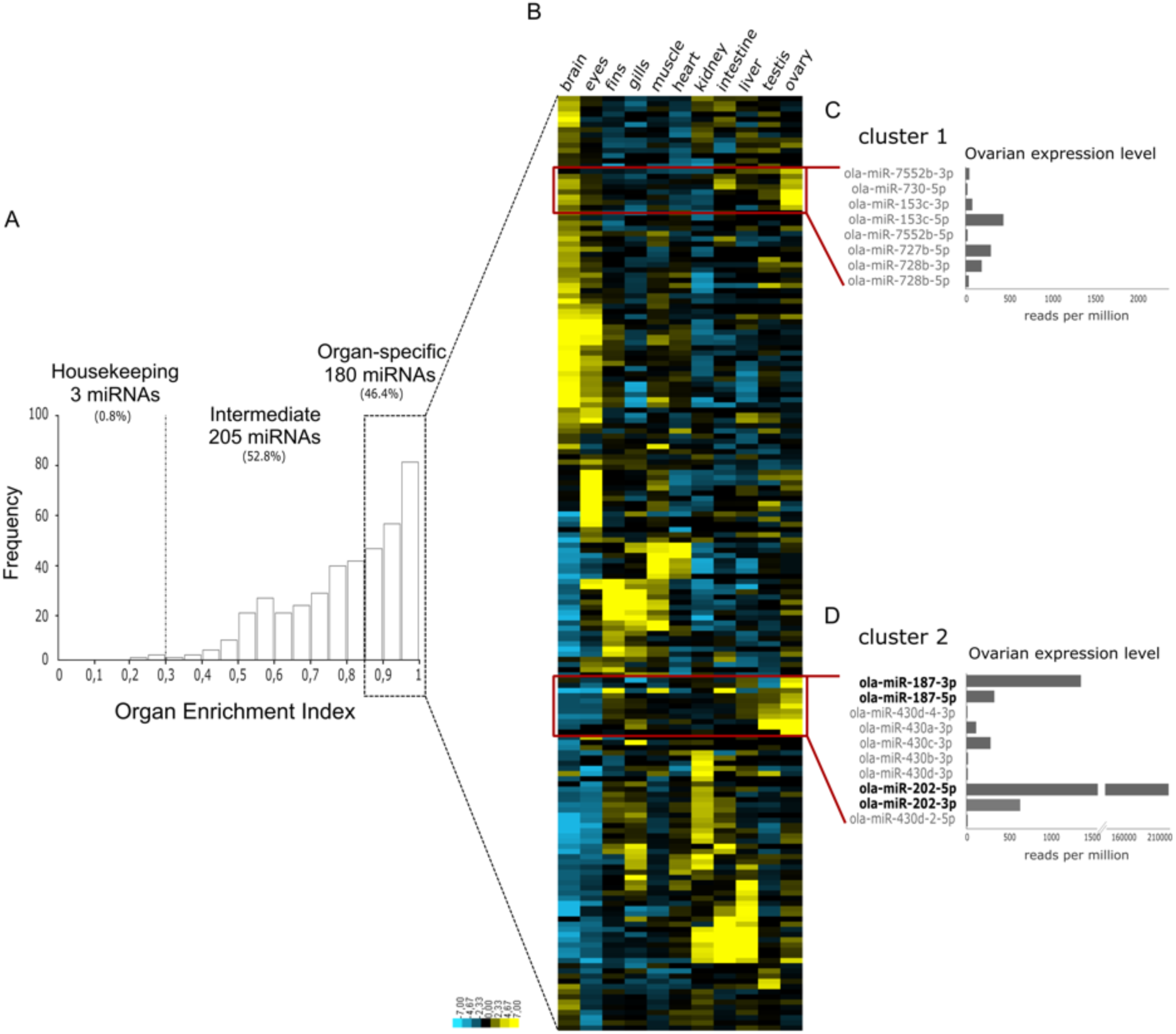
Ovarian enriched miRNAs. (A) Frequency distribution of Organ Enrichment Index (OEI) for 388 miRNAs across adult medaka tissues, including brain, eyes, fins, gills, muscle, heart, kidney, intestine, liver, testis, and ovary. (B) Heatmap of the 180 miRNAs (rows) enriched in at least one organ (columns). Expression values were median-centered prior to unsupervised hierarchical clustering using average linkage. Relative ovarian expression of ovarian-enriched miRNAs from cluster 1 (C) and cluster 2 (D). Color code: yellow indicates over-expression, blue indicates under-expression.

### *Mir-187* expression in ovary and brain

We quantified the expression levels of the two miR-187 mature forms (miR-187-5p and miR-187-3p) across adult tissues using reverse transcription quantitative polymerase chain reaction (TaqMan RT-qPCR, Fig 2). Both strands were detected exclusively in the ovary and the brain, with miR-187-3p exhibiting higher expression levels than miR-187-5p. Absolute expression levels (normalized to total RNA) were higher in the brain than in the ovary. However, their abundance relative to the total miRNA pool was lower in the brain (see Fig 1), consistent with the globally higher number of total miRNAs in brain tissue.

**Fig 2.**
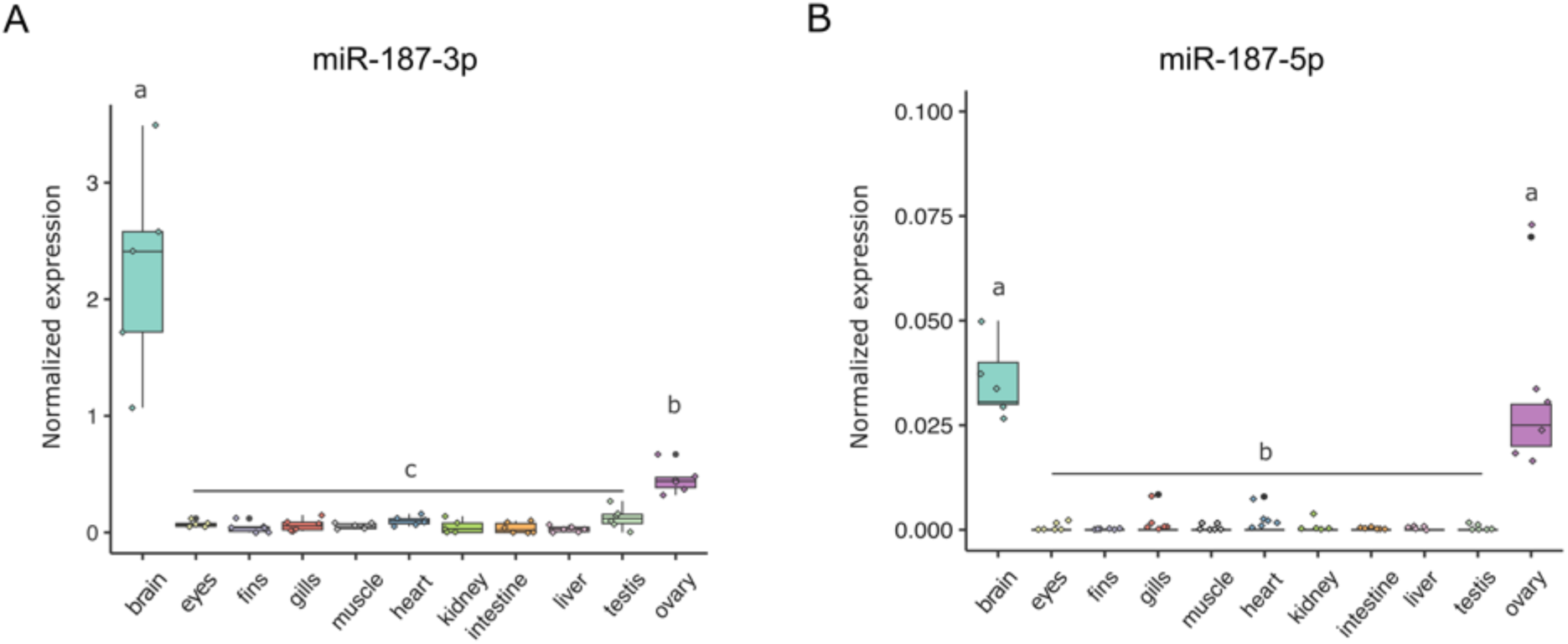
Expression levels of miR-187-3p and miR-187-5p in adult medaka tissues. Tissue distribution of miR-187-3p (A) and miR-187-5p (B) obtained by TaqMan RT-qPCR (n=5 or 6). Box plot edges define the 25th and 75th percentiles, the central line indicates the median, and whiskers show the 5th and 95th percentiles. Cel-miR-39-3p was used as an external calibrator for normalization to total RNA. Different letters indicate statistically significant differences between tissues, as determined by the non-parametric Kruskal-Wallis test followed by Dunn’s post hoc test (*p < 0.05*).

We then analyzed the spatial distribution of the predominant miR-187-3p mature form, using a single molecule *in situ* hybridization (smISH) assay (miRNAscope) on sections of ovary and brain from actively reproductive adult females, at 243 days post-fertilization (dpf) and 187 dpf, respectively. In the ovary, miR-187-3p was detected in oocytes at both pre-vitellogenic and vitellogenic stages (Fig 3A, A’, A’’), as well as in the granulosa cells surrounding developing follicles (Fig 3A’”, arrows). In the brain, miR-187-3p was detected in the dorsal hypothalamus (Fig 4B, B’), in cells surrounding the mesencephalic ventricle (Fig 4C, C’) and in Purkinje cells of the cerebellum (Fig 4D, D’). In both ovary and brain, no detectable signal above background was observed with the scramble negative control probe (Fig 3B and 4B”-4D”).

**Fig 3.**
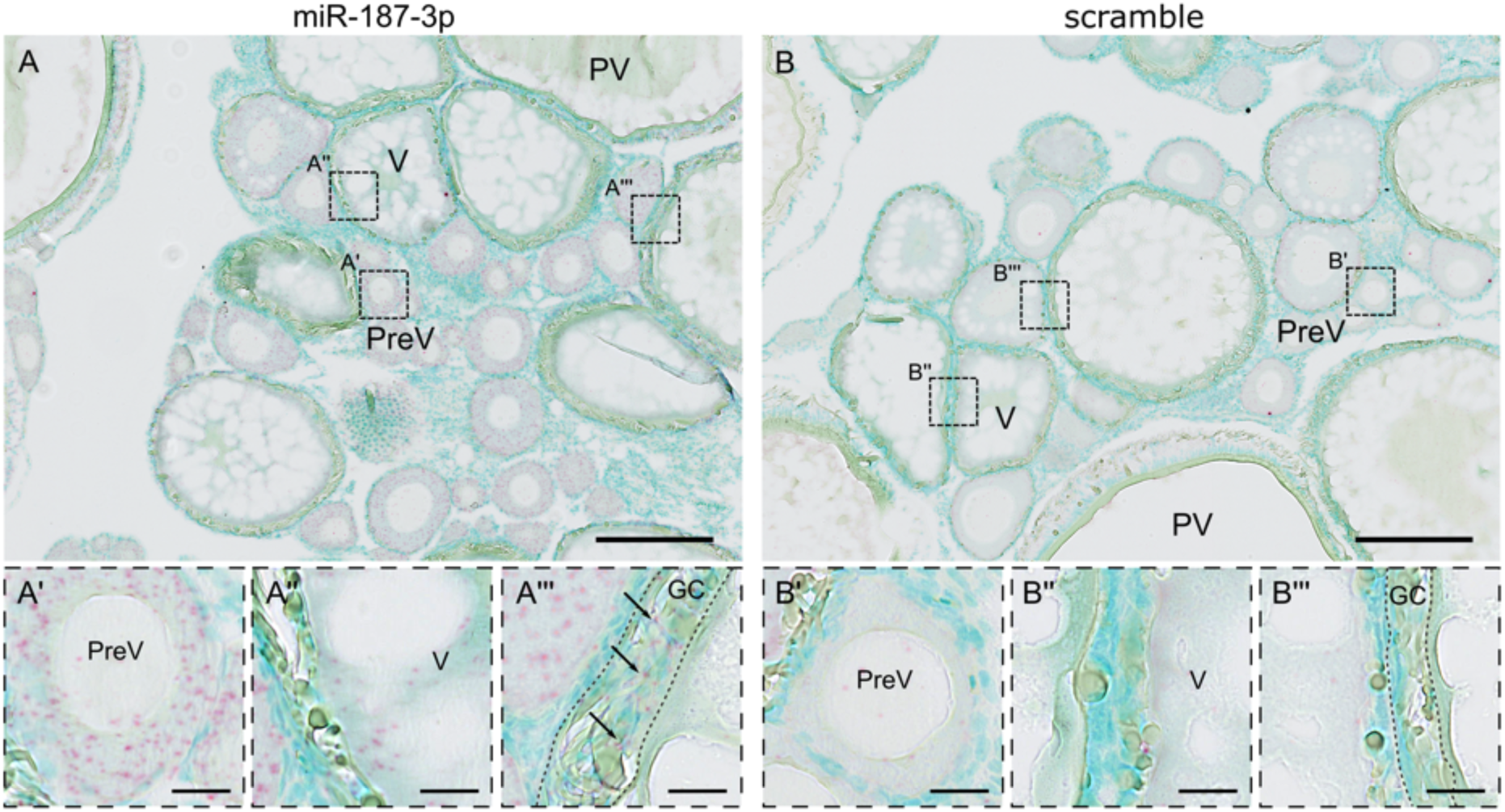
Expression pattern of miR-187-3p in the adult ovary. SmISH was performed on ovarian sections from adult female medaka at 243 dpf. (A) miR-187-3p was detected using specific miRNAscope probes (in red). Nuclei are stained with methyl green (in blue). (B) Negative scramble control probe. (A’ and B’) Higher magnification views of pre-vitellogenic oocytes. (A’’ and B”) Higher magnification views of vitellogenic oocytes. (A’’’ and B’’’) Higher magnification views of granulosa cells of a follicle during vitellogenesis. MiR-187-3p is detected in all oocytes of pre-vitellogenic follicles (from 25 to 150 µm diameter), in some vitellogenic follicles (up to ∼250 µm diameter), and in granulosa cells surrounding follicles at all stages. GCs: granulosa cells; PreV: pre-vitellogenic follicle; V: vitellogenic follicle; PV: post-vitellogenic follicle. Scale bars: 200 μm (A and B) and 20 μm (A’, A’’, A’’’, B’, B’’, B’’’).

**Fig 4.**
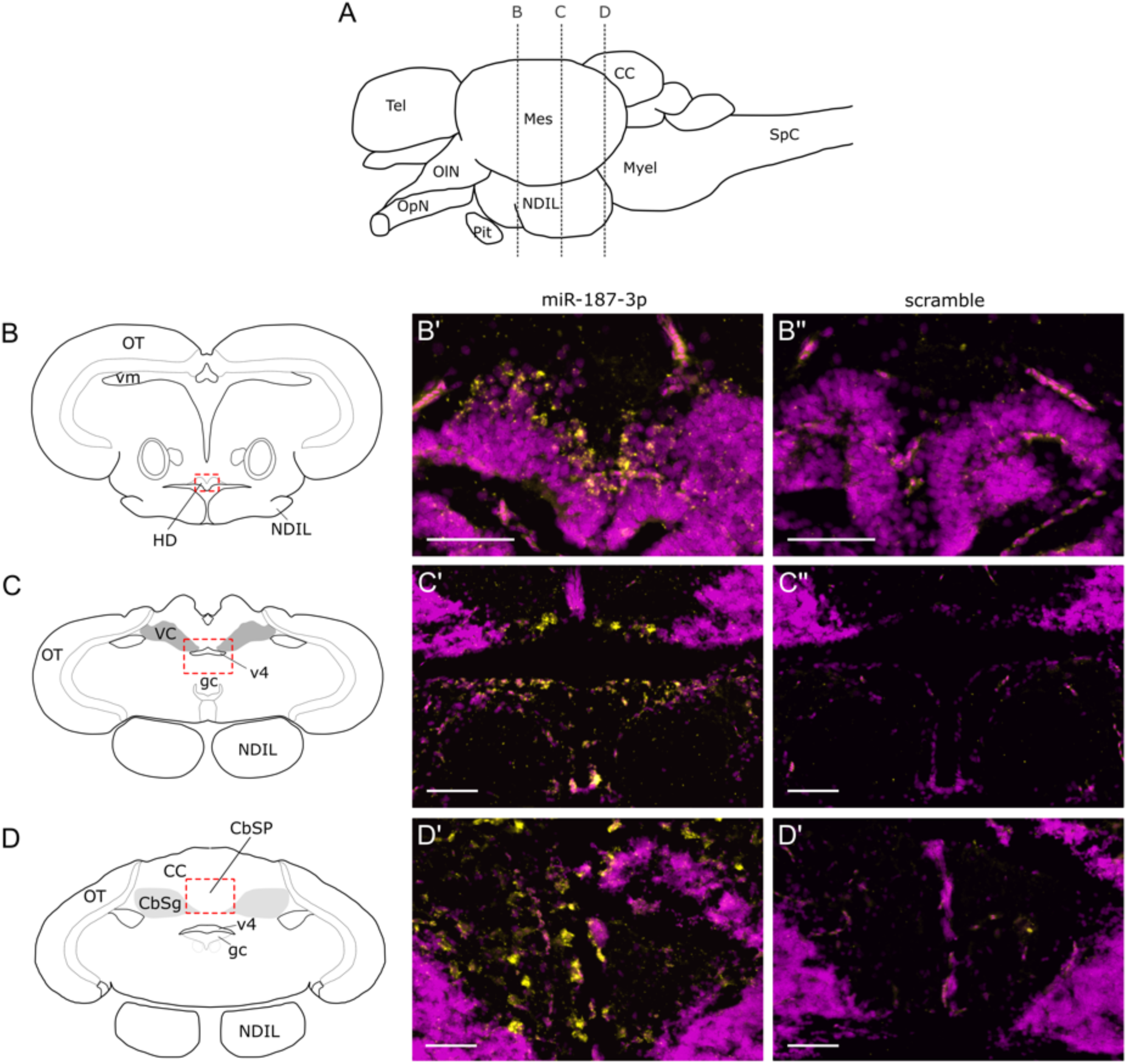
Expression of miR-187-3p in the adult medaka brain. SmISH was performed on brain sections from adult female medaka at 187 dpf. (A) Schematic illustration of the adult brain showing the positions of the transverse sections. Anterior is to the left, dorsal is at the top. (B-D) Diagrams of the transverse sections. (B’-D’) MiR-187-3p detection (in yellow). Nuclei are stained with methyl green (magenta). (B”-D”) Negative scramble control probe. miR-187-3p is expressed in cells of the dorsal hypothalamus (B’), in cells surrounding the mesencephalic ventricle (C’), and in Purkinje cells of the cerebellum (D’). Mes: mesencephalon, Tel: telencephalon, CC: corpus cerebelli, Myel: myelencephalon, SpC: spinal cord, OIN: olfactory nerve, OpN: optic nerve, NDIL: nucleus diffusus of lobus inferioris (hypothalamus), Pit: pituitary, OT: optic tectum, VC: valvula cerebelli, vm: ventriculus mesencephali, HD: hypothalamus periventricularis dorsalis, v4: ventriculus mesencephali, gc: griseum centrale, CbSg: stratum granulare of corpus cerebelli, CbSP: stratum ganglionare (Purkinje) of corpus cerebelli. Scale bars: 50 µm (B’-D’ and B’’-D”).

### *Mir-187* knock-out impairs female fecundity

To determine the role of *mir-187* in female reproduction, CRISPR/Cas9-mediated KO medaka line was generated. A 4 bp INDEL was introduced within the genomic region overlapping the PAM site and the 5’ arm of the precursor (Fig 5A). Hairpin secondary structure prediction indicated that this INDEL disrupted the canonical stem-loop conformation of the pri-miR-187 transcript (Fig 5B). In homozygous *mir-187* −/− mutant (MUT) females, the expression level of miR-187-3p in ovary, determined by TaqMan RT-qPCR, was nearly undetectable compared to *mir-187* +/+ wild-type (WT) females, and reduced by half in heterozygous *mir-187* +/− mutant (HET) females (Fig 5C). This indicated that the 4 pb INDEL was sufficient to abolish the miR-187-3p biogenesis in the KO line. The reproductive performance of MUT females was followed over 160 days after transfer to breeding long-day (LD) photoperiod regime, by monitoring daily egg production (*i.e.*, fecundity phenotype). No change in fertilization rate or embryonic developmental rate was observed. From approximately 40 days post-transfer (dpt) to the LD-photoperiod regime, the number of eggs laid over time by MUT females peaked at ∼ 15 eggs/day and remained consistently lower than for WT females, which reached ∼ 20 eggs/day (Fig 5D). This difference was also observed over the entire monitoring period, with a significantly reduced mean daily egg production for MUT compared to WT (12.8 ± 10.7 *vs.* 17.4 ± 11.7 eggs per day, respectively; Fig 5E). At the individual level, MUT females also displayed, over the period, a significantly lower average daily egg production than WT females (12.69 ± 6.04 *vs.* 18.73 ± 7.93 eggs per day per female, respectively Fig 5F).

**Fig 5.**
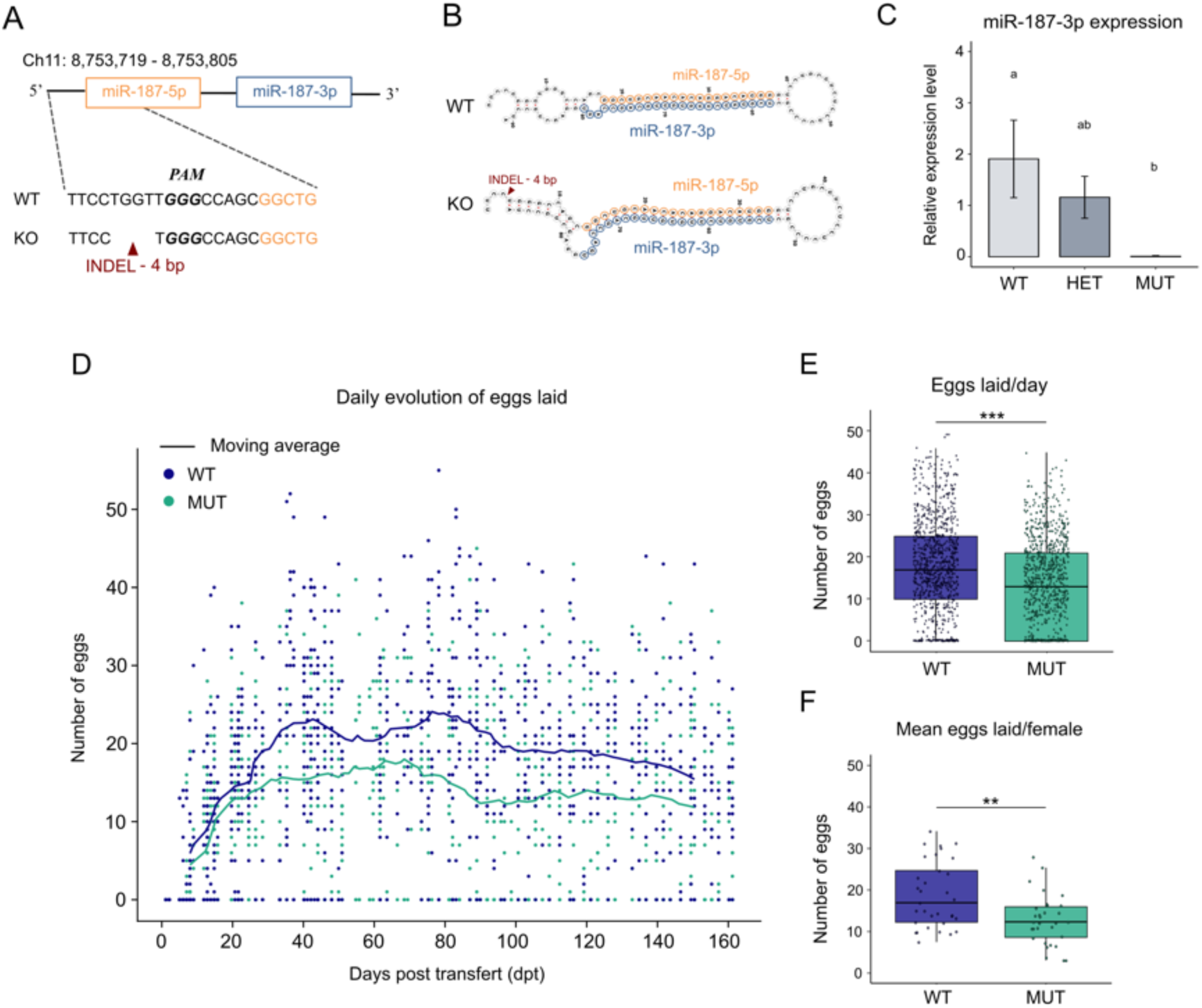
Generation of *mir-187* KO medaka line and reproductive phenotype. The *mir-187* gene was knocked out using CRISPR/Cas9 genome editing. Eggs laid by individual adult WT and MUT females were collected and counted for 160 days under a reproductive LD-photoperiod, and across three generations. (A) Schematic representation of the -4 bp INDEL mutation located upstream of the miR-187-5p sequence. The targeted PAM sequence is shown in bold. (B) Predicted hairpin secondary structure of pre-miR-187 for WT and KO fish. (C) Expression levels of mature miR-187-3p in ovaries from WT (*mir-187 +/+*), HET (*mir-187 +/-*) and MUT (*mir-187 -/-*) females at ∼30 dpf, measured by TaqMan RT-qPCR (n=3 per condition). Cel-miR-39-3p was used as an external control for normalization. Different letters indicate statistically significant differences between groups (Kruskal-Wallis test with Dunn’s post hoc test). (D) Daily evolution of egg laid during the spawning period. Solid lines represent the moving average over 11 days. The y-axis indicates the number of eggs laid per day; each dot represents one female’s egg production on a given day. DPF: days post fertilization; DPT: days post transfer. (E) Number of eggs laid per day. (F) Mean number of eggs laid each day per female. Boxplots show the interquartile range (25th-75th percentiles), with the median as a solid line. Whiskers extend to the 5th and 95th percentiles. Statistical significance was assessed using the Mann-Whitney test. Asterisks indicate significance levels: ** *p < 0.01*, *** *p < 0.001*.

### *Mir-187* knock-out induces a follicle growth delay

To further characterize the role of *mir-187*, we used 3D imaging to perform a spatially resolved cellular profiling of WT and MUT ovaries at 104 dpf under short-day (SD) photoperiod (non-spawning adult), and at 211 dpf under LD-photoperiod (spawning adult). Follicles of whole ovaries were quantified and categorized into developmental stages based on their diameter, according to the classification established by Iwamatsu *et al.* (Fig 6A, D)(20). At both ages, the total number of follicles per ovary was similar between genotypes, indicating that *mir-187* inactivation does not affect the overall size of the follicular pool (S1 Fig). When analyzing follicular diameter distributions at 104 dpf (non-spawning adult), we observed that MUT ovaries displayed a clear accumulation of stage I follicles (20-60 µm), and relatively fewer follicles at stages II (60-90 µm) and III (90-120 µm), compared to WT ovaries (Fig 6C, D and S1 Fig). This observation indicated a delay in early follicular growth in MUT ovaries, characterized by an accumulation of follicles at the earliest stage and a reduced transition to subsequent growth stages. At 211 dpf (spawning adult), follicles at all developmental stages (from stage I to stage IX) were present, consistent with the onset of full reproduction competence (S1 Fig) and no significant differences were detected between follicle stage distributions (Fig 6E-F). However, the density curves of the MUT group showed a subtle left shift compared to WT, with a mild accumulation of stage II/III follicles, and a moderately reduced density of later stage follicles compared to WT (Fig 6E and S1 Fig). Although not statistically significant, these trends suggest a possible mild but persistent delay in follicular progression in reproductively active females.

**Fig 6:**
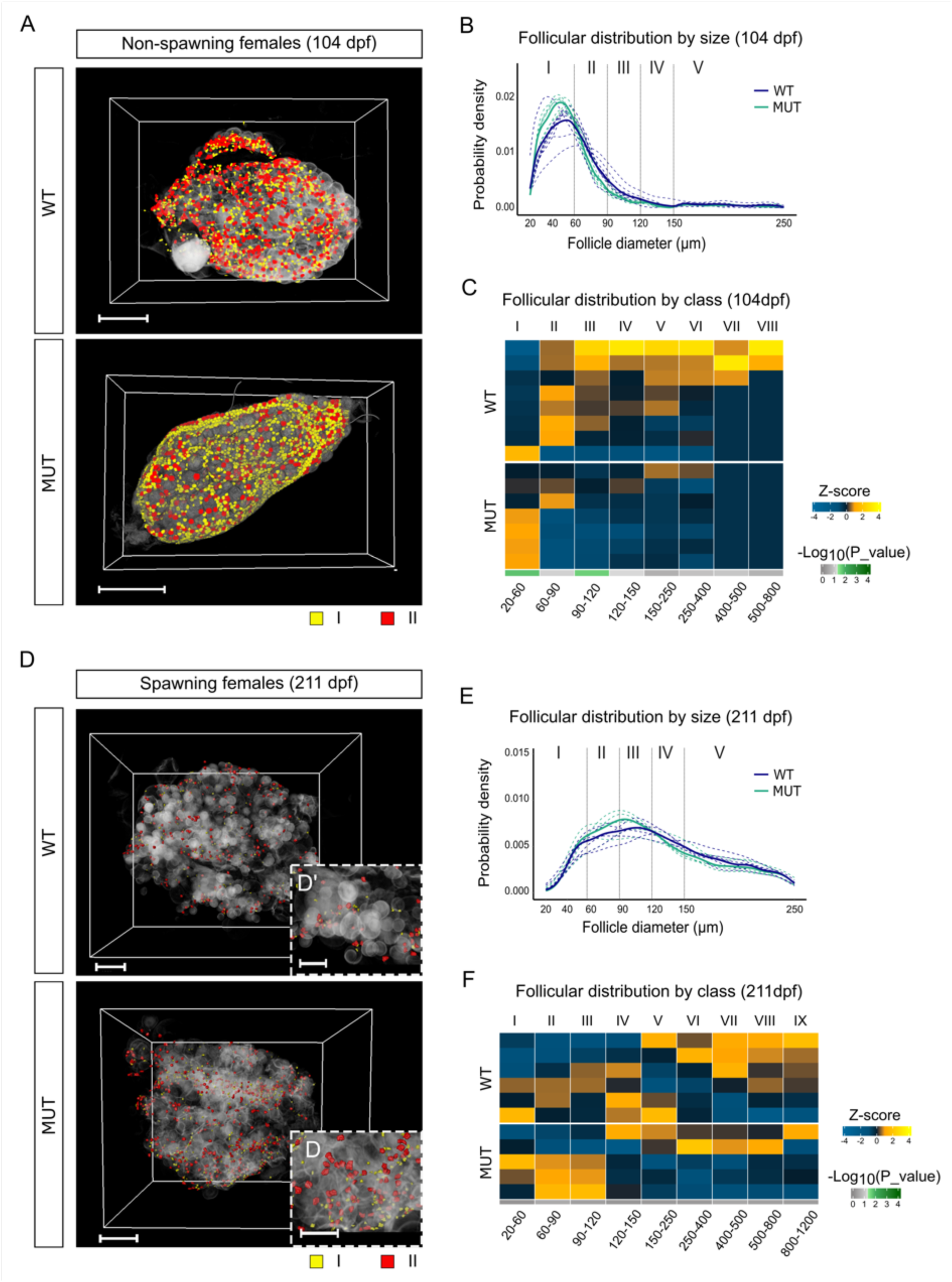
Follicular composition of WT and MUT ovaries from non-spawning and spawning females. Whole ovaries from non-spawning females at 104 dpf (under SD-photoperiod) and from spawning females at 211 dpf (under LD-photoperiod) were imaged in 3D using light-sheet fluorescence microscopy. 3D representation of WT (A) and MUT (D) ovaries at 104 dpf and 211 dpf, respectively, with segmented follicles at stage I (yellow) and II (red) highlighted. Scale bar: 1000 µm. Insets show zoomed views of stage I-II; scale bar: 500 µm. (B, E) Size probability density curves of 20−250 µm diameter follicles in WT and MUT at 104 dpf and 211 dpf. Solid lines show average follicle density; dotted lines represent individual samples. (C, F) Heatmaps representing relative follicular stage distribution in individual ovaries at 104 dpf and 211 dpf. For each ovary, the percentage of follicles per stage was converted into a z-score relative to the population mean. At 104 dpf, only stages I-VIII are shown, as no stage IX follicles were detected. Color intensity reflects relative abundance (warmer colors: higher z-score; cooler colors: lower z-scores). Horizontal bar below each heatmap represents Mann-Whitney test results per stage: green indicates statistically significance (p<0.05), grey tones indicate non-significant differences.

### *Mir-187* inactivation leads to molecular changes associated with follicle development

To investigate the molecular changes in *mir-187* KO, we performed a differential transcriptomic analysis of ovaries from WT and MUT females at 104 dpf (when stage I follicle accumulation is detectable in MUT), using Bulk RNA Barcoding and Sequencing (BRB-seq). Principal component analysis (PCA) revealed a clear separation between WT and MUT samples along the first principal component (PC1), which accounted for 98.16% of the variance (Fig 7A). A total of 2,884 differentially expressed genes (DEGs; adjusted *p*-value < 0.05) were identified, with WT and MUT samples forming two distinct clusters (Fig 7B). KEGG pathway enrichment analysis of the 1,459 most significantly deregulated DEGs (0.5 < log₂ fold change < -0.5) showed significant enrichment for pathways related to hormonal signaling (*i.e.*, “cAMP signaling pathway”, “estrogen pathways”, …), cell proliferation and differentiation (*i.e.*, “signaling pathway regulating pluripotency of stem cells”), and cell adhesion (e.g., “focal adhesion” and “cytoskeleton in muscle cells”, Fig 7C and S2 Fig). These results illustrate a broad transcriptomic remodeling of the ovary in *mir-187* KO fish at 104 dpf.

**Fig 7.**
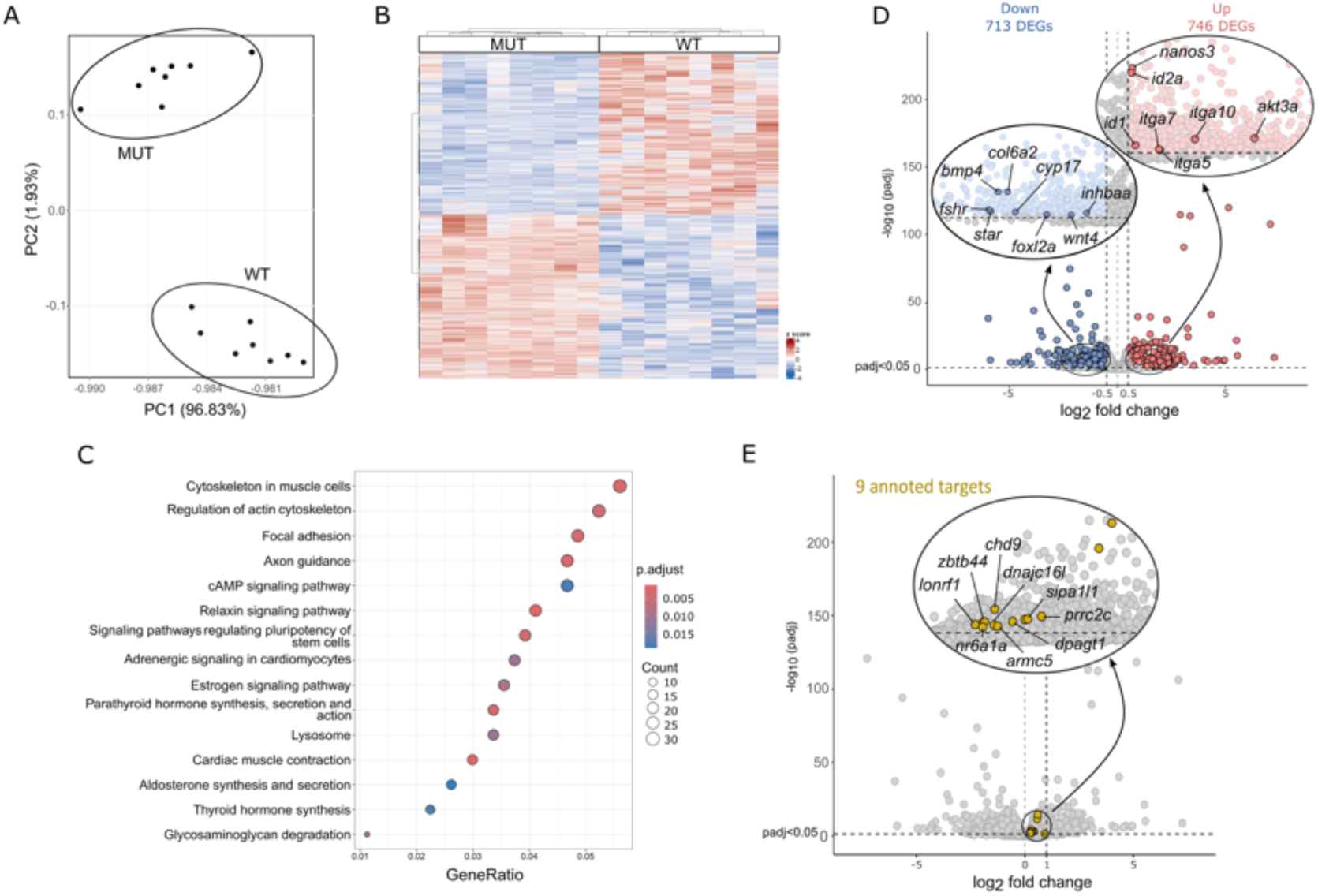
Transcriptomic analysis and possible targets of miR-187-3p in ovary. Comparison between WT and MUT medaka ovaries from young non-spawning adults (104 dpf, n=8 per group). (A) Principal Component Analysis (PCA) of WT and MUT samples, with each point representing one biological replicate. PC1 (x-axis) accounts for 98.16% of the variance, and PC2 (y-axis) for 0.74%. Group separation along PC1 indicates major transcriptomic differences between WT and MUT groups. (B) Heatmap illustrating the expression profiles of 2,884 DEGs identified between WT and MUT samples. Expression values were scaled per gene (z-score) to highlight relative expression differences across samples (red: higher expression, blue: lower expression). Hierarchical clustering of samples (top dendrogram) reflects transcriptomic similarities. Rows represent genes and columns represent individual samples. (C) Dot plots showing dysregulated KEGG molecular signaling pathways, based on enrichment of 1.459 most differentially expressed genes (log_2_FC<-0.5 or log_2_FC >0.5). This enrichment was calculated using identified human orthologs mapped to the KEGG signaling pathway database. Dot size indicates the number of DEGs involved in each pathway, and color represents the adjusted p-value (more significant in red). The y-axis lists pathways names, and the x-axis shows the proportion of DEGs mapped to each pathway. (D) Volcano plot displaying 1.459 DEGs in ovaries of MUT *vs* WT medaka, selected based on a log_2_ fold change threshold of <-0.5 and >0.5. Among them, 713 DEGs are significantly down-regulated in MUT (blue) and 746 are up-regulated (red). A zoomed-in view emphasizes several genes with known roles in oogenesis. (E) Identification of potential phenotypic targets of miR-187-3p in ovaries. Selected genes were filtered based on an *in silico* prediction of a highly active target site (TargetScan and 7-8 mer), with a low fold-change in up-regulation (0<log2FC<1), inter-individual variability limits (IVV_WT_<FC(MUT/WT)), and a higher coefficient of variation in MUT than WT (CV_MUT_>1.1×CV_WT_). A total of 14 possible targets in the ovary, 9 of which are annotated (zoomed-in view).

Among the 1,459 DEGs, we observed the down-regulation of several key genes with well-established roles in follicle development and growth (Fig 7D). Specifically, we identified down-regulation of genes encoding key steroidogenic genes (*fshr*, *star* and *cyp17*), genes involved in granulosa cell differentiation and follicle development, including a key effector of the canonical Wnt/β-catenin signaling pathway (*wnt4a*), forkhead transcription factors (*foxl2a*, *foxo1a* and *foxo3b*), as well as genes of the TGF-β signaling pathway (*inhbaa*, *acvr1b* and *bmp4*), all of which being essential for granulosa cell differentiation and steroid hormone biosynthesis. Down-regulation of these genes indicates an impaired steroidogenic program and follicle developmental pathways in MUT ovaries.

Among the 1,459 DEGs, we also observed the up-regulation of several genes associated with early follicle activation and follicular cell proliferation (Fig 7D). Notably, the kinase *akt3a*, a component of the PI3K/Akt pathway, showed increased expression, consistent with the concomitant down-regulation of its downstream FOXO targets (foxo1a, foxo3b) previously described. The inhibitors of differentiation *id1* and *id2a*, which are commonly induced in growth-factor–responsive contexts, were also markedly upregulated. In addition, the germ-cell marker *nanos3*, required for the maintenance of early oocyte identity, showed increased expression in MUT. Together, these transcriptional changes point to enhanced proliferative or maintenance signals within both somatic and germ-cell compartments, suggesting a shift toward a more undifferentiated cellular state at early stages of follicle development.

Finally, we observed the dysregulation of several genes involved in cell adhesion and extracellular matrix (ECM) organization in MUT ovaries (Fig 7D). Analysis of the pathways involved in cell adhesion, revealed that a large number of genes in the collagen family, such as *col1a1b, col1a2, col6a3, col6a2* and *col4a1*, were down-regulated. These collagens are essential structural components of the extracellular matrix (ECM) (21). In contrast, several integrins, such as *itga7, itga10* and *itga5*, were up-regulated. Integrins are transmembrane receptors that mediate cell adhesion to the ECM and activate intracellular signaling cascades (22). This dysregulation points to a remodeling of the ECM and ovarian tissue integrity in MUT ovaries, which may impair mechanical support, cellular communication, and finally compromise follicular development and fertility.

### MiR-187-3p candidate target genes

Using BRB-seq data, we searched for candidate functional targets of miR-187-3p among the numerous putative target genes (*i.e.*, up-regulated genes containing a predicted miR-187-3p binding site in their 3′ UTR). We applied an efficient pipeline to highlight relevant possible miRNA target genes (Janati-Idrissi et al., 2024). This screening strategy is based on two main criteria, including limited fold-change and a low inter-individual variability in gene expression in miRNA KO samples. We identified 9 annotated genes (*nr6a1a*, *prrc2c*, *lonrf1*, *dpagt1*, *sipa1l1*, *chd9*, *dnajc16l*, *armc5,* and *zbtb44*) (Fig 7E and Fig S3). Among them, two targets (*nr6a1a* and *dpagt1*) have previously been implicated in follicular development. Interestingly, *nr6a1a* (also known as germ cell nuclear factor, GCNF), is a nuclear receptor that represses pluripotency genes and regulates oocyte-specific transcriptional programs.

## DISCUSSION

MicroRNAs are increasingly recognized as key regulators of ovarian development and reproductive function in vertebrates, including teleost fish (3–5). However, most functional insights still come from *in vitro* studies, and *in vivo* evidence remains limited in fish. Here, we identify *mir-187* as one of the most ovarian-enriched miRNA in medaka, second only to *mir-202* (23). We show that the predominant (*i.e.*, biologically active) mature form miR-187-3p in the ovary is expressed in both oocytes and granulosa cells, and that *mir-187* loss-of-function disrupts follicle developmental dynamics and reduces female fecundity. While these results clearly demonstrate that mir-187-3p acts within the ovary to modulate oogenesis, a brain-mediated contribution cannot be entirely excluded, as mir-187-3p was found to be also expressed in discrete brain territories, including cerebellar Purkinje cells (24) and the dorsomedial hypothalamus (25).

Integrated analyses combining high-resolution 3D imaging and transcriptomic profiling revealed that the reduced fecundity of *mir-187* MUT most likely results from a deficit in the recruitment and subsequent progression of early previtellogenic follicles (stage I). 3D reconstructions of ovaries from non-spawning adults showed an accumulation of stage I follicles accompanied by a reduced proportion of more advanced stages. This pattern reflects a delay in follicle progression limiting the number of oocytes that reach maturation and ovulation, rather than a global developmental delay of the fish, or a modification in the overall follicle pool size, both of which remain unchanged. This delay is accompanied by a transcriptional signature indicating impaired follicular growth, paracrine signaling and steroidogenesis. In particular, down-regulation of regulators of the Wnt/β-catenin pathway (wnt4a) (26), of the TGF-β pathway (inhbaa, acvr1b, bmp4) (27–29), and of several key steroidogenic genes (fshr, star, cyp17) (30–33). Globally, such transcriptomic perturbations suggest defective somatic cell proliferation and insufficient support for follicle growth, which is consistent with the delay observed by 3D imaging in the early transition of stage I follicles to more advanced stages.

In mammals, the PI3K/Akt–FOXO pathway is an important regulator of the early steps of primordial follicle activation, and similar roles have been proposed in teleost (34). In medaka, this activation step likely corresponds to the exit of the early primordial follicles (*i.e.*, follicles arrested in the diplotene cell-cycle stage, Gdip) from the germinal cradle and their entry into newly formed primary follicles (*i.e.*, follicles at stage I)(35). In mammals, the PI3K/Akt pathway contributes to this process notably through the phosphorylation and inactivation of FOXO transcription factors, which are key regulators of follicular quiescence and somatic homeostasis (36,37). In *mir-187* KO medaka ovaries, the up-regulation of *akt3a* together with down-regulation of *foxo1a* and *foxo3b* strongly suggests a dysregulation of this conserved pathway at the time of early follicle activation. Although direct evidence for PI3K/Akt control of the Gdip-to-stage I transition is still lacking in medaka, this transcriptional changes in the *mir-187* MUT strongly support a model in which the earliest activation step could be abnormally regulated. In parallel, the marked up-regulation of *id1* and *id2a*, two inhibitors of somatic differentiation commonly induced by TGF-β/BMP-responsive pathways, suggests that granulosa cells remain in an immature state (38,39). Together, impaired FOXO signaling and sustained Id expression likely suggest that the differentiation of somatic cells is impaired, thereby promoting premature activation of primordial follicles (Gdip) while preventing their proper progression beyond stage I. This dual mechanism could provide a coherent hypothesis to explain the accumulation of early stage I follicles observed in the *mir-187* MUT.

In addition to follicle developmental defects, *mir-187* KO led to a marked upregulation of *nanos3*, an evolutionarily conserved germline stem cell gene essential for the development and maintenance of primordial germ cells in fish (40–42). This suggests that the defects observed in *mir-187* null fish might also arise prior to follicle assembly, within the germinal cradle itself, highlighting the need to investigate potential alterations in the proliferation and/or specification of oogonia cells that could also contribute to the accumulation of stage I follicles.

Regarding the post-transcriptional regulatory mechanisms of miR-187-3p, we identified 7 candidate targets (*nr6a1a*, *prrc2c*, *lonrf1*, *dpagt1*, *sipa1l1*, *chd9*, and *zbtb44*). Only two of them have known ovarian function. The first one (*dpagt)* encodes the enzyme catalyzing the first step of protein *N*-glycosylation, and has been shown to be essential for oocyte development and female fertility in mice (43). The second one, the *gcnf* gene (Germ Cell Nuclear Factor, also known as *nr6a1a*), is of particular interest. In mouse, GCNF contributes to regulating paracrine communication between the oocyte and somatic cells by repressing the expression of key TGFβ signaling components, *BMP-15* and *GDF-9*, thereby regulating early follicle development (44). In medaka, *gcnf* is specifically expressed in early oocytes and is therefore well positioned to act during the initial germ cell-oocyte transition (45). However, its role in fish ovary remains unknown, and no downstream targets have yet been characterized. The identification of *gcnf* as a putative miR-187-3p target thus provides a compelling entry point to explore a previously unrecognized regulatory pathway acting at the onset of folliculogenesis in teleost. Determining whether *gcnf* fulfills a conserved role in medaka, potentially through modulation of BMP/GDF signaling, and validating its regulation by miR-187-3p through targeted knockout, will thus be of utmost interest for future studies.

Identifying direct targets of miRNAs is notoriously challenging, as neither computational predictions nor *in vitro* assays provide information on the functional relevance of individual interactions (7,46). Nevertheless, two putative targets have been proposed for miR-187 based such approaches in mammals, namely the *tgfbr2* gene in pigs and the *bmpr2* gene in bovine cumulus cells (47,48). In our study, we applied a dedicated prediction pipeline integrating differential expression data (13), however, none of these genes emerged as candidates targets of miR-187-3p. While this discrepancy may reflect species-specific target repertoires, it more likely highlights the methodological difficulties inherent to miRNA target identification. Undoubtedly, it will be necessary to determine which of these genes, when deregulated, reproduce the observed phenotypic perturbations. This will require *in vivo* functional validation using target-site mutants, notably by disrupting the miR-187-3p binding site within the 3′ UTR of candidate genes and assessing whether this disruption recapitulates the *mir-187* KO phenotype (7,49).

Finally, the reproductive phenotype of *mir-187* KO females (*i.e.*, reduction in egg production) shows partial overlap with that of *mir-202*, one of the few microRNAs functionally characterized in the medaka ovary (14). However, *mir-202* KO females display a much stronger reduction in female fecundity and an additional defect in the quality of eggs produced (*i.e.*, eggs that cannot be fertilized), consistent with its direct action on *tead3b* and its role in early follicular growth and granulosa cell differentiation (13). In contrast, we observed that the milder *mir-187* phenotype involves dysregulation of genes controlling early follicle activation and somatic maturation, and that it is associated with distinct predicted target genes, pointing to a regulatory pathway distinct from that of *mir-202*. These differences suggest that the two miRNAs act at different steps of early folliculogenesis and through non-overlapping target networks. Determining whether their functions converge or operate independently will require further dedicated analyses.

In summary, we shed light on a novel actor in oogenesis regulation and provide the first *in vivo* evidence that *mir-187* controls early follicle development and female fecundity in medaka. We showed that loss of *mir-187* leads to the accumulation of stage I follicles likely through combined defects in follicle activation and somatic cells maturation. Together with *mir-202*, *mir-187* is now one of the few microRNAs demonstrated to contribute to medaka female fecundity, and more generally to reproductive success in fish, with a seemingly earlier and mechanistically distinct role in controlling follicle recruitment. Our findings thus broaden the landscape of microRNA-mediated regulation in fish oogenesis.

## MATERIAL AND METHODS

### Ethics statement

Fish were reared and handled in the INRAE LPGP fish facility, which holds full approval for animal experimentation (C35-238-6). All experiments were conducted in compliance with the European directive 2010/63/EU on the protection of animals used for scientific purposes and were approved by the INRAE-LPGP Institutional Animal Care and Use Committee under #M-2021-13-VT-SG [14-VT-SG], #M-VTWTFW-4222, and #M-VT002F8-4522 [F8-5222, F10-4623].

### Small RNAseq data analysis

Small RNAseq data available on the NCBI Sequence Read Archive under accession number SRP151190 (14), and in the FishmiRNA database (www.fishmirna.org)(10), were analyzed. Raw reads were processed using Prost! Software version 0.7.60. (17). MiRNAs between 17 and 25 bp, and with a total rpm count exceeding 10, were kept. A scalar quantitative measure of miRNA expression specificity across different tissues was calculated according to the original index (tau) developed by Yanai et al. (18). An organ enrichment index (OEI) was calculated for a given miRNA j as follows:

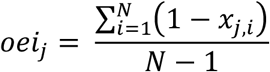

where *N* is the total number of tissues, and *x_i,j_* represents the expression intensity of tissue *i*, normalized by the tissue with the highest expression for miRNA *j*. The resulting values range from 0 to 1. The closer the value is to 0, the more ubiquitously the miRNA is expressed; conversely, the closer the value is to 1, the more tissue-specific the miRNA is. A clustering algorithm was applied to the most tissue-specific miRNAs (OEI>0.85) to identify ovarian miRNA-enriched clusters. All data were centered and normalized to a median of zero. This unsupervised clustering analysis was performed using CLUSTER and TREEVIEW software.

### Fish breeding

Juvenile and adult female medaka fish (*Oryzias latipes*) from the CAB strain were raised at 26°C under controlled photoperiod conditions. Fish were raised under a short-day (SD-photoperiod regime (12 h light/12 h dark), at a maximum density of 25 fish in 10 L tanks, until approximately 104 dpf. Then fish were transferred to a long-day LD-photoperiod regime (15 h 45 light/8 h 15 dark) that triggers reproduction.

### Reproductive performance

Each female (WT or MUT) was paired with a WT male in a 1.4 L tank, and individual clutch monitoring was performed from the beginning of the reproductive phase, after transfer to LD-photoperiod regime, at 104 dpf (0 dpt), up to 270 dpf (160 dpt). The number (*i.e.*, fecundity) and development (*i.e.*, quality) of eggs laid was recorded daily. To assess the fecundity phenotype, a moving average over 11 time points was applied to smooth individual trajectories. This phenotyping approach was conducted across three independent experiments, each involving different generations of medaka females, to ensure the robustness and reproducibility of the observations. For a given day *t_j_*, the mean number of eggs laid was determined by averaging the values for that day, along with the six previous and six subsequent data points, as follows:

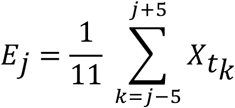

where *X_ti_* is the mean number of eggs laid on day *t_i_*. This computed average value was assigned to the mean abscissa of the 11 data points:

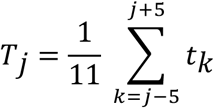

### Establishment of the *mir-187* KO medaka line

For CRISPR/Cas9 mediated knock-out experiment, a short target sequence with the 20bp-NGG motif was identified in the *mir-187* gene with the help of the CRISPOR online tool (http://crispor.tefor.net/) and the medaka genome (UCSC Genome Browser on Medaka Oct. 2005, NIG/UT MEDAKA1/oryLat2 Assembly). A specific miR-187 guide (miR-187-sgRNA) was constructed using the pDR274 vector (Plasmid 42250, Addgene) and the One-Step Overlap-PCR method (OSO-PCR)(50). The OSO forward primer (5’-gTGCAGCCGCTGGCCCAACC-GTTTTAGAGCTAGAAATAGCAAG-3’) and its reverse complement, OSO reverse primer (5’-gGTTGGGCCAGCGGCTGCAC-TATAGTGAGTCGTATTAGC-3’) were added in limiting amounts to a PCR reaction with external primers: universal primer forward (5’-CAGGAAACAGCTATGAC-3’) and universal primer reverse (5’-AAAAGCACCGACTCGGTG-3’). PCR was performed with the JumpStart^TM^ Taq Polymerase (D9307, Sigma) under the following conditions: 94°C for 5 min; and 38 cycles of 94°C for 30 sec, 55°C for 30 sec, and 72°C for 3 min. The T7-target-sgRNA PCR product (262bp) was excised on an agarose gel and purified using the NucleoSpin^TM^ Gel and PCR Clean-up kit (740609, Macherey Nagel), and was then used for *in vitro* transcription of the sgRNA using a MEGAshortscript^TM^ T7 transcription kit (AM1354, Life Technologies). The Cas9-RNA was transcribed from the pCS2-nCas9n vector (Plasmid 47929, Addgene), linearized with NotI and using the mMESSAGE mMACHINE^TM^ SP6 transcription Kit (AM1340, Life Technologies). Both Cas9-RNA and sgRNA were purified using a standard phenol/chloroform protocol. Cas9-RNA (100 ng.μl^-1^) and miR-187-sgRNA (10 ng.μl^-1^) were co-injected into one-cell stage embryos. Injected embryos were reared to sexual maturity and INDEL mutations were by genotyping. Founder fishes (F0) carrying the same small 4-nucleotide deletion upstream of the mature miR-187-5p sequence were backcrossed with WT F0 fish (*mir-187+/+*) to produce HET F1 fish (*mir-187+/-*) and establish a stable mutant line. HET F1 fish were intercrossed to generate homozygous WT and MUT offspring, and females were kept for phenotypic analyses.

### Genotyping

#### Genomic DNA extraction

For INDEL detection and genotypic sex determination, genomic DNA (gDNA) was extracted from a small piece of the caudal fin sampled from adult fish anesthetized with tricaine at 225 mg/L (PharmaQ) diluted in water with 450 mg/L of sodium bicarbonate (Sigma S5761). Fin samples were lysed in 60 µl of lysis buffer containing 1.25 M NaOH and 10 mM EDTA (pH 12), incubated at 90°C for 1 h, and then neutralized with 60 µL of neutralization solution containing 2 M Tris-HCl (pH 5).

#### PCR for sex determination

Genotypic sex was determined by PCR amplification on gDNA targeting the *dmrt1* gene. In medaka, *dmrt1a* (autosomal, located on chromosome 9), and *dmrt1Y* (Y-linked paralog) can be distinguished. The primer pair (Forward: AGTGCTCCCGCTGCCGGAACCAC, Reverse: AATTCTGGCATCTTTGCAGTCAGCA) amplifies both *dmrt1a* and *dmrt1Y*. PCR was performed in 25 μl reactions containing 2.5 µl of gDNA diluted (1:10), 2.5 µl 10x buffer, 0.25 µl dNTPs at 10 mM, 0.5 µl each primer at 10 µM, and 0.25 µl Jumpstart polymerase (Sigma, D9307). Products were separated by electrophoresis on a 1.5% agarose gel in 1X TAE buffer and stained with SYBR Safe. Female individuals showed a single band (*dmrt1a*), while male individuals exhibited two bands (*dmrt1a* and *dmrt1Y*).

#### PCR for INDEL detection

To identify WT and mutant (INDEL -4) alleles, the gDNA was amplified by qPCR around the expected mutation site, using a common reverse primer (miR-187_R: GGTCCCAGCACTCTCATGTG) and two allele-specific forward primers: one for the WT allele (miR-187-WT_F: CAGCCGCTGGCCCAACCA), and one for the MUT allele (miR-187-Mut_F: CAGCCGCTGGCCCAGGA). Quantitative PCR was performed in 10 μl reactions containing 4 µl of gDNA diluted (1:10), 0.5 µl each primer at 10 µM, and 5 µl PowerUp SYBR Green Master Mix polymerase (Applied Biosystems, A25742), following the manufacturer’s protocol. Quantitative PCR was carried out on a StepOnePlus thermocycler (Applied Biosystems) with the following program: 95 °C for 3 minutes for initial denaturation, followed by 40 cycles of 95 °C for 3 sec, and 60 °C for 30 sec.

### Tissues collection

For tissues/organs dissection, fish were euthanized by immersion in a lethal dose of tricaine (PharmaQ, 300 mg/L) supplemented with sodium bicarbonate (600 mg/L, Sigma S5761). For TaqMan RT-qPCR and BRB-seq, collected samples were immediately frozen in liquid nitrogen and subsequently stored at −80°C until RNA extraction. For miRNAscope and 3D imaging, all samples were first dissected and then fixed in 4% paraformaldehyde (PFA) in 1x PBS (overnight at 4 °C followed by 4 hours at RT), except for ovaries, which were dissected in PBS after fixation in PFA. All samples were then gradually dehydrated in 100% MeOH, and stored at −20°C until use.

### Total RNA extraction

Total RNA was extracted from collected tissues for TaqMan RT-qPCR and BRB-seq. Frozen tissues were lysed with Precellys Evolution Homogenizer (Ozyme, Bertin technologies) in TRIS Reagent (TR118, Euromedex) and total RNA was extracted according to manufacturer’s instructions. RNA quality was checked using an Agilent 2100 Bioanalyzer, and RIN values were ≥ 8 for all samples. RNA quantity was measured using a NanoDrop^TM^ 1000 Spectrophotometer (Thermo Scientific, USA).

### TaqMan quantitative RT-PCR

For reverse transcription (RT), 60 ng of total RNA and 60 fmol.µl^-1^ of an external calibrator cel-miR-39-3p (478293_miR, Life technologies) were used. The TaqMan advanced miRNA cDNA Synthesis Kit was used according to manufacturer’s instructions (A28007, Applied Biosystems). Quantitative PCR was then performed using 2.5 μl of diluted cDNA (1:5), 0.5 μl of TaqMan Advanced miRNA Assay solution (CCU001S, Special Product Custom designed advanced miRNA assay, Life technologies) and 5 μl of Fast Advanced Master Mix (4444557, Applied Biosystems) in a total volume of 10 μl. The LightCycler 480 system (Roche) was used under the following conditions: 95°C for 20 sec, followed by 40 cycles of 95°C for 1 sec and 60°C for 20 sec. MiRNA expression levels were calculated from a standard curve using LightCycler^®^ 480 software (Roche). Samples were run in duplicate, and the standard curve in triplicate. The external miRNA cel-miR-39-3p was used for normalization.

### BRB-seq experiment

Bulk RNA barcoding and sequencing (BRB-seq) was performed on total RNA extracts. Libraries were constructed using the Mercurius BRB-seq kit (PN 10813, v0.6.2. revA, Nov. 2023) from Alithea Genomics. Clustering and sequencing were performed on an Illumina NovaSeq 6000 using the SBS (Sequencing By Synthesis) technology with the NovaSeq Reagent Kits. Samples were sequenced on an SP flow cell in paired-end 28x90 mode. Over 0.6 billion reads were assigned, including 1 to 3 million reads for brain and 3.5 to 18 million reads for ovary.

### Differential expression analysis

Genome alignment and exon-level quantification was carried out with BRB-seq reads using STARsolo (51). These steps have been integrated in the form of a Nextflow RNAseq pipeline available on GitLab (https://forge.inrae.fr/lpgp/rnaseq) and archived on Zenodo (DOI 10.5281/zenodo.15260130). Genes (pseudo) counts (>= 10 reads per gene in at least 4 samples per condition), normalization, and differential expression comparisons across groups were carried out using the DESeq2 package (v1.42.1) for each comparison, to prevent large observed higher within-group gene variability observed, which can inflate the per-gene dispersion estimate for other group comparisons (52). DEGs were first identified using an adjusted p-value threshold of padj < 0.05. To refine the selection and highlight the most strongly deregulated genes, an additional filtering criterion was applied, retaining only those with a log_2_ fold change greater or less than 0.5.

### KEGG pathway enrichment

Differentially expressed genes obtained with BRB-seq were mapped to their human orthologs using the getBM() function from the biomaRt package (v2.58.2). KEGG pathways enrichment was performed based on human orthologs (https://www.genome.jp/kegg/pathway.html) using the enrichKEGG() function (organism = “hsa”) from the KEGGREST package (v1.42.0). Pathways associated with the “Human Diseases” category were intentionally excluded from the enrichment results. Adjusted p-values were computed using the Benjamini-Hochberg procedure, and pathways with an adjusted p-value < 0.05 were considered significantly enriched.

### MicroRNA folding and target identification

The sequence of miR-187-3p was used for *in silico* prediction of the hairpin secondary structure, based on minimum free energy structures and base pair probabilities, using the RNAfold web server with default settings (http://rna.tbi.univie.ac.at/cgi-bin/RNAWebSuite/RNAfold.cgi)(53). For the identification of phenotypic miR-187-3p targets, we followed the Janati-Idrissi *et al*. pipeline (13) that applies successive filtering steps to TargetScan-predicted targets (54). First, only 7-mer and 8-mer seed-matched targets were retained. Second, based on BRB-seq data, only targets showing moderate overexpression in the MUT compared to WT were kept (0 < log2FC < 1). Third, targets with high inter-individual variability in WT relative to MUT were excluded (IVV_WT_ < FC(MUT/WT)). Finally, only targets displaying a higher coefficient of variation in MUT than in WT were retained (CV_MUT_ > 1.1 × CV_WT_).

### miRNAscope

Brains and ovaries collected from fishes, at 187 and 243 dpf, respectively, were embedded in paraffin, and sectioned at 7 μm thickness using a microtome (microm, HM355). Superfrost Plus slides (VWR) were used to ensure better adhesion of the sections during the staining procedure. The staining was performed with the miRNAscope HD Reagent Kit-RED (ACDbio, 324500) according the manufacturer’s instructions (ACDBio, Bio-techne, Abingdon, United Kingdom). Probes targeting specific miRNAs were designed and synthesized by ACDBio. The following probes were used: SR-ola-miR-202-5p-S1 (1127801-S1), SR-ola-miR-202-3p-S1 (1127811-S1), SR-ola-miR-187-3p-O1-S1 (1127821-S1), along with a negative control probe SR-Scramble-S1 (727881-S1). Design of the ola-miR-187-5p probe was not possible due to the high risk of cross-hybridization with endogenous transcripts such as *tsr2*, *rbm19* and *clock*. Signal detection was achieved by incubating the tissue sections with FastRED-B, diluted 1:60 in FastRED-A for 15 min at RT in the dark. Nuclear staining was performed using methyl green (MG, 40 µg.ml^-1^) for 1 hour at RT protected from light. After a brief xylene wash and air drying, coverslips were mounted with EcoMount mounting medium (ACDBio, 320409). Whole-slide imaging was performed using a NanoZoomer slide scanner equipped with a 40x objective lens (Hamamatsu).

### 3D imaging

Ovaries collected from WT and MUT females at 104 dpf (*i.e.*, non-reproductively active females in SD-photoperiod regime) and at 211 dpf (*i.e.*, reproductively active females in LD-photoperiod regime), were treated following the cubic−ethyl cinnamate combined clearing method (C-ECi) with a few modification (55). Nuclear staining was performed by incubating ovaries with the MG fluorescent dye (40 μg.ml^-1^, 323829, Sigma-Aldrich) diluted in PBS/0.1% Triton X-100 at room temperature (RT) for 2.5 days. After staining, ovaries from 211 dpf females were thoroughly rinsed in PBS and then embedded in 1.2% standard agarose blocks to facilitate mounting for subsequent light-sheet microscopy imaging. All samples were subsequently dehydrated through a graded series of methanol/H2O dilutions and cleared by immersion in 100% ECi (W243000, Sigma-Aldrich), and stored at RT until imaging. Ovaries from 104 dpf females were imaged with a Leica TCS SP8 laser scanning confocal microscope equipped with a 16x/0.6 IMM CORR VISIR HC FLUOTAR objective (15506533, Leica), optimized for immersion acquisition mode in ECi. Images were acquired with a 638 nm laser excitation for MG, in 512×512 pixels, 400 Hz (bidirectional), optical zoom 0.75, z-steps 3 μm (voxel size 1.8×1.8×3 μm), line accumulation and frame average set to 2. For complete confocal imaging, samples were successively placed with ventral side facing upwards and then downwards, due to the limited working distance of the objective, as described previously (55). Ovaries from 211 dpf females were imaged using an UltraMicroscope Blaze^TM^ lightsheet microscope (Miltenyi Biotec, Bergisch Gladbach, Germany) equipped with a 4x objective lens (NA 0.35), enabling rapid acquisition of large samples. Samples were placed in a plastic holder from Miltenyi, secured with a screw to maintain consistent orientation throughout the acquisition process, placed in a tank filled with an organic solvent (MACS Imaging Solution, Miltenyi Biotec, RI 1.556–1.562), and illuminated by the laser light sheet from both sides left and right (light sheet thickness 3.91µm). Images were acquired with a laser 640 nm for MG, a camera sCMOS (exposure time 50ms; 2048X2048), and a z-step of 4 µm (voxel size: 1.62x1.62x4 µm). All datasets were acquired at equal imaging parameters (exposure time, numerical aperture, sheet width).

### Image analysis

3D image analysis for follicle segmentation in whole ovaries (including image preprocessing, deep learning−based segmentation, postprocessing, and diameter measurement) was performed using Cellpose 2.2.2 (Python 3.10.16)(56), FIJI (57) and Amira 3D software (version 2020.2, https://www.thermofisher.com/amira), following the pipeline previously described by Lesage *et al*., with minor modifications (58). An additional step was introduced to facilitate the removal of false-positive labels generated during Cellpose segmentation, particularly in images of 211 dpf adult ovaries. These artifacts were primarily caused by the agarose surrounding the ovaries during light-sheet acquisition. A custom Cellpose model was trained to generate a mask outlining the entire ovary. The resulting mask was inverted to create a background label mask, which was then applied to the images to eliminate background-associated false-positive labels.

## ACKNOWLEDGMENTS

We thank the INRAE-LPGP fish facility staff, and especially Brigitte Guillet, Cécile Duret, and Guillaume Gourmelen, for fish rearing and husbandry. We are grateful to the GenoToul bioinformatics platform Toulouse Occitanie (Bioinfo GenoToul, doi: 10.15454/1.5572369328961167E12) for providing support with computing and storage resources. We thank the MGX-Montpellier GenomiX platform for BRB-sequencing. We thank the APEX imaging facility (INRAE Oniris, UMR703 PAnTher, labeled IBiSA and Biogenouest, member of the NeurATRISinfrastructure) for access to the light-sheet microscope, and the LPGP 3D imaging facility for access to the confocal microscope. We also thank the University of Rennes 1 H2P2 facility for the use of the slide scanning NanoZoomer. We are grateful to Constance Merdrignac for helpful discussions and for providing scripts and guidance on KEGG pathway analysis.

## ADDITIONAL INFORMATION AND DECLARATIONS

### Authors Contributions

M.D. and S.G.: Investigation, Methodology, Formal analysis, Resources, Writing – original draft.

M.T., S.H., F.M., L.D., J.M., A.B.: Methodology, Software, Visualization.

J.B.: Conceptualization. Writing – review & editing.

V.T.: Conceptualization, Formal analysis, Funding acquisition, Investigation, Project administration, Supervision, Writing – review & editing.

All authors approved the final version.

### Data Availability

All BRBseq datasets, 3D images, image labels, and image-derived diameters generated in this study are publicly available. BRBseq datasets are available on the NCBI SRA portal under accession PRJNA1233206. The 3D images and their corresponding image labels are available on the BioImage Archive repository under DOI 10.6019/S-BIAD1824. The 3D image-derived diameters are available on Recherche Data Gouv (https://doi.org/10.57745/B6OOMX). Scripts used to generate 3D image labels, compute 3D image-derived diameters, and perform BRBseq data analysis are available on GitHub (https://github.com/VThermes-LPGP) and archived on Zenodo under DOIs (10.5281/zenodo.17989576 and 10.528/zenodo.17989255).

**S1 Fig:**
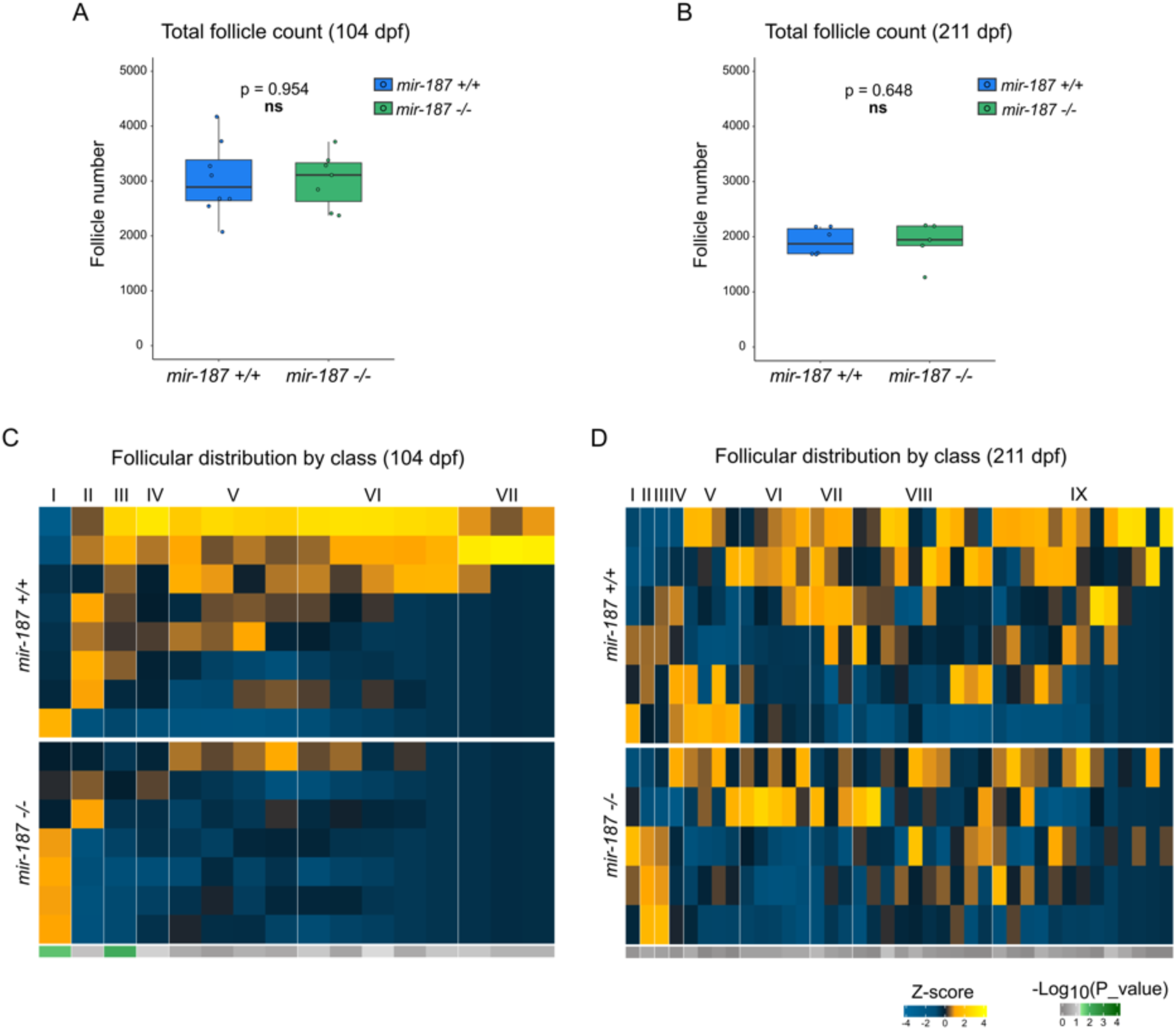
Follicular total count and distribution of WT and MUT ovaries at 104 and 211 dpf. Ovaries from young adults under short-day photoperiod (104 dpf) and adults under long day photoperiod (211 dpf) were imaged using light-sheet fluorescence microscopy. (A, C) Total number of follicles per ovary in *mir-187+/+* and *mir-187-/-* females at 104 dpf (A) and 211 dpf (C). Each dot represents a single ovary. Boxplots show medians and interquartile ranges; whiskers indicate the 5^th^ and 95^th^ percentiles. Statistical comparisons were using Mann-Whitney tests; p-value are indicated on the plots. (B, D) High resolution heatmaps showing the relative distribution of follicles in *mir-187+/+* and *mir-187-/-* ovaries at 104 dpf (B) and 211 dpf (D), using 30µm intervals. This fine-grained resolution improves the visualization of size variations between early and later follicular stages. Each row represents an individual ovary; color intensity reflects z-score values (warmer colors: above average; cooler colors: below average). A horizontal bar below each heatmap shows results of Mann-Whitney tests per stage: green indicates statistically difference (*p<0.05*), grey tones indicate non-significant differences.

**S2 Fig:**
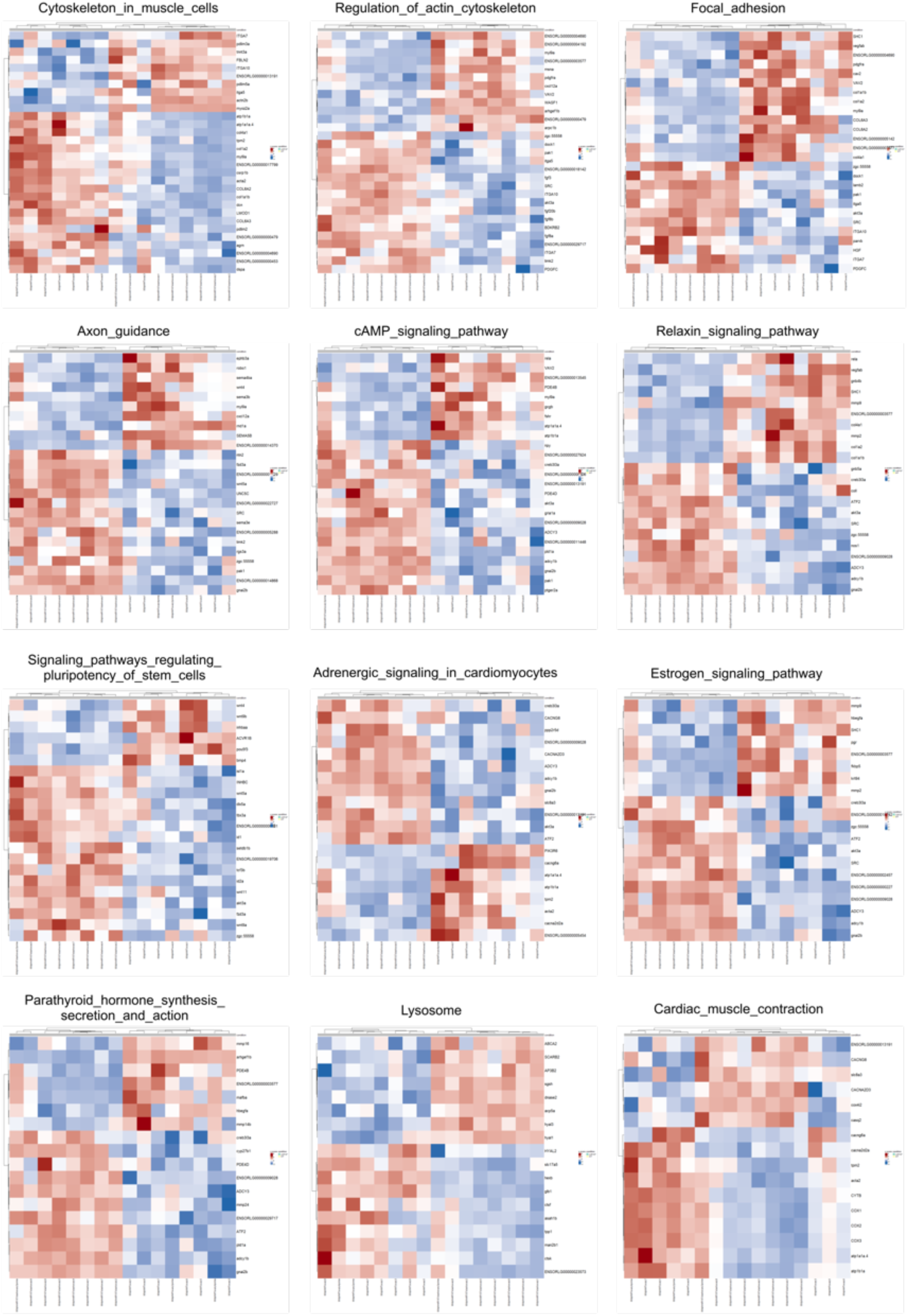
Heatmaps of differentially expressed pathways in *mir-187* MUT and WT medaka ovaries. Heatmaps of selected KEGG pathways significantly enriched in the ovarian transcriptome. Rows show pathway genes and columns represent Mut and WT replicates. Expression values are Z-score scaled per gene (red: upregulated; blue: downregulated). Differential regulation is observed in pathways related to cytoskeletal organization, signal transduction, hormone signaling, and muscle contraction.

**S3 Fig:**
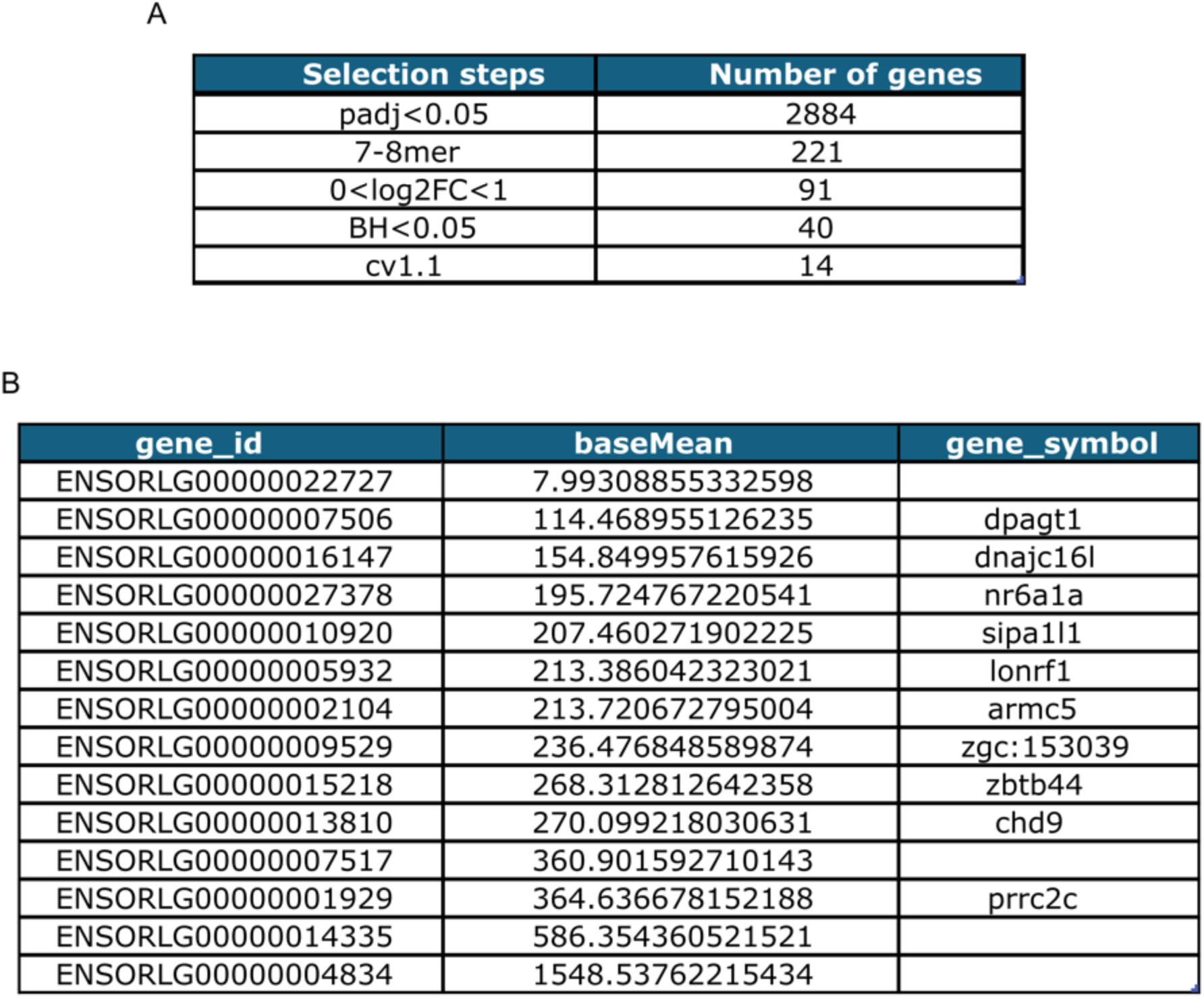
Selection of predicted miR-187-3p target genes. (A) Number of predicted miR-187-3p target genes remaining after each filtering step, including statistical significance (padj<0.05), seed type (7-8mer), fold-change(0<log2FC<1), and variability criteria (IVV_WT_<FC(MUT/WT) or BH<0.05 and CV_MUT_>1.1×CV_WT_). (B) Final list of 14 selected genes after applying all selection filters, with their mean expression values and gene symbols.

## REFERENCES

1. Iwamatsu T, Ohta T, Oshima E, Sakap N. Oogenesis in the Medaka Oryzias latipes-Stages of Oocyte Development. Unpublished. 1988;

2. Lubzens E, Young G, Bobe J, Cerdà J. Oogenesis in teleosts: How fish eggs are formed. General and Comparative Endocrinology. 2010;165:367–89.

3. Bizuayehu TT, Babiak I. MicroRNA in teleost fish. Genome Biology and Evolution. 2014;6:1911–37.

4. Baley J, Li J. MicroRNAs and ovarian function. J Ovarian Res [Internet]. 2012 [cited 2021 Nov 30];5(1):8. Available from: http://ovarianresearch.biomedcentral.com/articles/10.1186/1757-2215-5-8

5. Alvi SM, Zayed Y, Malik R, Peng C. The emerging role of microRNAs in fish ovary: A mini review. General and Comparative Endocrinology. 2021;311:113850.

6. Lai EC. Two decades of miRNA biology: lessons and challenges. RNA [Internet]. 2015 Apr [cited 2018 Mar 7];21(4):675–7. Available from: https://www.ncbi.nlm.nih.gov/pmc/articles/PMC4371328/

7. Bartel DP. Metazoan MicroRNAs. Cell [Internet]. 2018 Mar [cited 2021 Nov 30];173(1):20–51. Available from: https://linkinghub.elsevier.com/retrieve/pii/S0092867418302861

8. Ambros V. MicroRNA-mediated gene regulation and the resilience of multicellular animals. Postepy Biochem [Internet]. 2024 May 23 [cited 2026 Jan 7];70(1):62–70. Available from: https://postepybiochemii.ptbioch.edu.pl/index.php/PB/article/view/515

9. Seitz H. A new perspective on microRNA-guided gene regulation specificity, and its potential generalization to transcription factors and RNA-binding proteins. Nucleic Acids Research [Internet]. 2024 Sept 9 [cited 2025 Dec 17];52(16):9360–8. Available from: https://academic.oup.com/nar/article/52/16/9360/7734163

10. Desvignes T, Bardou P, Montfort J, Sydes J, Guyomar C, George S, et al. FishmiRNA: An Evolutionarily Supported MicroRNA Annotation and Expression Database for Ray-Finned Fishes. Molecular Biology and Evolution. 2022;39:1–7.

11. Xiao J, Zhong H, Zhou Y, Yu F, Gao Y, Luo Y, et al. Identification and Characterization of MicroRNAs in Ovary and Testis of Nile Tilapia (Oreochromis niloticus) by Using Solexa Sequencing Technology. PLoS One [Internet]. 2014 Jan 23 [cited 2018 Mar 9];9(1). Available from: https://www.ncbi.nlm.nih.gov/pmc/articles/PMC3900680/

12. Donadeu FX, Schauer SN, Sontakke SD. Involvement of miRNAs in ovarian follicular and luteal development. Journal of Endocrinology [Internet]. 2012 Dec [cited 2026 Jan 6];215(3):323–34. Available from: https://joe.bioscientifica.com/view/journals/joe/215/3/323.xml

13. Janati-Idrissi S, de Abreu MR, Guyomar C, de Mello F, Nguyen T, Mechkouri N, et al. Looking for a needle in a haystack: de novo phenotypic target identification reveals Hippo pathway-mediated miR-202 regulation of egg production. Nucleic Acids Research. 2024;52:738–54.

14. Gay S, Bugeon J, Bouchareb A, Henry L, Montfort J, Le Cam A, et al. MicroRNA-202 (miR-202) controls female fecundity by regulating medaka oogenesis. bioRxiv. 2018;287359.

15. Xiong S, Tian J, Ge S, Li Z, Long Z, Guo W, et al. The microRNA-200 cluster on chromosome 23 is required for oocyte maturation and ovulation in Zebrafish. Biology of Reproduction. 2020;103(4):769–78.

16. Ludwig N, Leidinger P, Becker K, Backes C, Fehlmann T, Pallasch C, et al. Distribution of miRNA expression across human tissues. Nucleic Acids Research. 2016;44(8):3865–77.

17. Desvignes T, Batzel P, Sydes J, Eames BF, Postlethwait JH. miRNA analysis with Prost! reveals evolutionary conservation of organ-enriched expression and post-transcriptional modifications in three-spined stickleback and zebrafish. Scientific Reports. 2019;9(1):1–15.

18. Yanai I, Benjamin H, Shmoish M, Chalifa-Caspi V, Shklar M, Ophir R, et al. Genome-wide midrange transcription profiles reveal expression level relationships in human tissue specification. Bioinformatics. 2005;21(5):650–9.

19. Tani S, Kusakabe R, Naruse K, Sakamoto H, Inoue K. Genomic organization and embryonic expression of miR-430 in medaka (Oryzias latipes): Insights into the post-transcriptional gene regulation in early development. Gene. 2010;449:41–9.

20. Iwamatsu T, Ohta T, Oshima’ E, Sakap N. Oogenesis in the Medaka Oryzias latipes-Stages of Oocyte Development. Vol. 5, ZOOLOGICAL SCIENCE. 1988.

21. Ricard-Blum S. The Collagen Family. Cold Spring Harbor Perspectives in Biology [Internet]. 2011 Jan 1 [cited 2025 July 18];3(1):a004978–a004978. Available from: http://cshperspectives.cshlp.org/lookup/doi/10.1101/cshperspect.a004978

22. Barczyk M, Carracedo S, Gullberg D. Integrins. Cell Tissue Res [Internet]. 2010 Jan [cited 2025 July 18];339(1):269–80. Available from: http://link.springer.com/10.1007/s00441-009-0834-6

23. Bouchareb A, Le Cam A, Montfort J, Gay S, Nguyen T, Bobe J, et al. Genome-wide identification of novel ovarian-predominant miRNAs: new insights from the medaka (Oryzias latipes). Sci Rep [Internet]. 2017 Jan 10 [cited 2017 Jan 27];7. Available from: http://www.ncbi.nlm.nih.gov/pmc/articles/PMC5223123/

24. Locke TM, Fujita H, Hunker A, Johanson SS, Darvas M, du Lac S, et al. Purkinje Cell-Specific Knockout of Tyrosine Hydroxylase Impairs Cognitive Behaviors. Frontiers in Cellular Neuroscience. 2020;14(July):1–16.

25. Gooley JJ, Schomer A, Saper CB. The dorsomedial hypothalamic nucleus is critical for the expression of food-entrainable circadian rhythms. Nature Neuroscience. 2006;9(3):398–407.

26. Boyer A, Lapointe É, Zheng X, Cowan RG, Li H, Quirk SM, et al. WNT4 is required for normal ovarian follicle development and female fertility. FASEB j [Internet]. 2010 Aug [cited 2025 July 18];24(8):3010–25. Available from: https://onlinelibrary.wiley.com/doi/abs/10.1096/fj.09-145789

27. Li CW, Zhou R, Ge W. Differential regulation of gonadotropin receptors by bone morphogenetic proteins in the zebrafish ovary. General and Comparative Endocrinology [Internet]. 2012 May [cited 2026 Jan 5];176(3):420–5. Available from: https://linkinghub.elsevier.com/retrieve/pii/S0016648011005260

28. Akimoto Y, Fujii W, Naito K, Sugiura K. The effect of ACVR1B/TGFBR1/ACVR1C signaling inhibition on oocyte and granulosa cell development during *in vitro* growth culture. J Reprod Dev [Internet]. 2023 [cited 2025 July 18];69(5):270–8. Available from: https://www.jstage.jst.go.jp/article/jrd/69/5/69_2023-041/_article

29. Zhao C, Zhai Y, Geng R, Wu K, Song W, Ai N, et al. Genetic analysis of activin/inhibin β subunits in zebrafish development and reproduction. Mullins MC, editor. PLoS Genet [Internet]. 2022 Dec 5 [cited 2025 July 18];18(12):e1010523. Available from: https://dx.plos.org/10.1371/journal.pgen.1010523

30. Lubzens E, Young G, Bobe J, Cerdà J. Oogenesis in teleosts: How fish eggs are formed. General and Comparative Endocrinology. 2010;165:367–89.

31. Stocco DM. StAR Protein and the Regulation of Steroid Hormone Biosynthesis. Annu Rev Physiol [Internet]. 2001 Mar [cited 2025 July 18];63(1):193–213. Available from: https://www.annualreviews.org/doi/10.1146/annurev.physiol.63.1.193

32. Kitano T, Takenaka T, Takagi H, Yoshiura Y, Kazeto Y, Hirai T, et al. Roles of Gonadotropin Receptors in Sexual Development of Medaka. Cells [Internet]. 2022 Jan 24 [cited 2025 July 18];11(3):387. Available from: https://www.mdpi.com/2073-4409/11/3/387

33. Rajendiran P, Jaafar F, Kar S, Sudhakumari C, Senthilkumaran B, Parhar IS. Sex Determination and Differentiation in Teleost: Roles of Genetics, Environment, and Brain. Biology [Internet]. 2021 Sept 27 [cited 2025 Nov 19];10(10):973. Available from: https://www.mdpi.com/2079-7737/10/10/973

34. Das D, Arur S. Conserved insulin signaling in the regulation of oocyte growth, development, and maturation. Mol Reprod Dev [Internet]. 2017 June [cited 2025 Nov 18];84(6):444–59. Available from: https://onlinelibrary.wiley.com/doi/10.1002/mrd.22806

35. Nakamura S, Kobayashi K, Nishimura T, Higashijima S, Tanaka M. Identification of Germline Stem Cells in the Ovary of the Teleost Medaka. Science. 2010;1561.

36. Shen M, Liu Z, Li B, Teng Y, Zhang J, Tang Y, et al. Involvement of FoxO1 in the effects of follicle-stimulating hormone on inhibition of apoptosis in mouse granulosa cells. Cell Death Dis [Internet]. 2014 Oct 16 [cited 2025 July 18];5(10):e1475–e1475. Available from: https://www.nature.com/articles/cddis2014400

37. Hu CL, Cowan RG, Harman RM, Quirk SM. Cell Cycle Progression and Activation of Akt Kinase Are Required for Insulin-Like Growth Factor I-Mediated Suppression of Apoptosis in Granulosa Cells. Molecular Endocrinology [Internet]. 2004 Feb 1 [cited 2025 July 18];18(2):326–38. Available from: https://academic.oup.com/mend/article/18/2/326/2747496

38. Su Y, Zheng L, Wang Q, Bao J, Cai Z, Liu A. The PI3K/Akt pathway upregulates Id1 and integrin α4 to enhance recruitment of human ovarian cancer endothelial progenitor cells. BMC Cancer [Internet]. 2010 Dec [cited 2025 July 18];10(1). Available from: https://bmccancer.biomedcentral.com/articles/10.1186/1471-2407-10-459

39. Hogg K, Etherington SL, Young JM, McNeilly AS, Duncan WC. Inhibitor of Differentiation (Id) Genes Are Expressed in the Steroidogenic Cells of the Ovine Ovary and Are Differentially Regulated by Members of the Transforming Growth Factor-β Family. Endocrinology [Internet]. 2010 Mar 1 [cited 2025 July 18];151(3):1247–56. Available from: https://academic.oup.com/endo/article/151/3/1247/2456622

40. Nishimura T, Fujimoto T. Generation of primordial germ cell-like cells by two germ plasm components, dnd1 and nanos3, in medaka (Oryzias latipes). iScience [Internet]. 2025 Mar [cited 2025 Nov 19];28(3):111977. Available from: https://linkinghub.elsevier.com/retrieve/pii/S2589004225002378

41. Köprunner M, Thisse C, Thisse B, Raz E. A zebrafish *nanos*-related gene is essential for the development of primordial germ cells. Genes Dev [Internet]. 2001 Nov 1 [cited 2025 Nov 19];15(21):2877–85. Available from: http://genesdev.cshlp.org/lookup/doi/10.1101/gad.212401

42. Aoki Y, Nakamura S, Ishikawa Y, Tanaka M. Expression and Syntenic Analyses of Four *nanos* Genes in Medaka. Zoological Science [Internet]. 2009 Feb [cited 2025 Nov 19];26(2):112–8. Available from: http://www.bioone.org/doi/abs/10.2108/zsj.26.112

43. Li H, You L, Tian Y, Guo J, Fang X, Zhou C, et al. DPAGT1-Mediated Protein *N*-Glycosylation Is Indispensable for Oocyte and Follicle Development in Mice. Advanced Science [Internet]. 2020 July [cited 2025 July 18];7(14). Available from: https://onlinelibrary.wiley.com/doi/10.1002/advs.202000531

44. Lan ZJ. GCNF-dependent repression of BMP-15 and GDF-9 mediates gamete regulation of female fertility. The EMBO Journal [Internet]. 2003 Aug 15 [cited 2025 July 18];22(16):4070–81. Available from: http://emboj.embopress.org/cgi/doi/10.1093/emboj/cdg405

45. Xie Z, Song P, Zhong Y, Guo J, Gui L, Li M. Medaka gcnf is a component of chromatoid body during spermiogenesis. Aquaculture and Fisheries [Internet]. 2021 Nov [cited 2025 July 18];6(6):574–82. Available from: https://linkinghub.elsevier.com/retrieve/pii/S2468550X20300745

46. Prajapat MK, Maria AG, Vidigal JA. CRISPR-based dissection of miRNA binding sites using isogenic cell lines is hampered by pervasive noise. Nucleic Acids Research [Internet]. 2025 Jan 7 [cited 2025 Dec 17];53(1):gkae1138. Available from: https://academic.oup.com/nar/article/doi/10.1093/nar/gkae1138/7915247

47. Fu Y, Zhang JB, Han DX, Wang HQ, Liu JB, Xiao Y, et al. CiRS-187 regulates BMPR2 expression by targeting miR-187 in bovine cumulus cells treated with BMP15 and GDF9. Theriogenology. 2023;197:62–70.

48. Yang L, Du X, Wang S, Lin C, Li Q, Li Q. A regulatory network controlling ovarian granulosa cell death. Cell Death Discovery. 2023;9:1–11.

49. Ambros V. MicroRNA-mediated gene regulation and the resilience of multicellular animals. Postepy Biochem [Internet]. 2024 May 23 [cited 2025 Dec 17];70(1):62–70. Available from: https://postepybiochemii.ptbioch.edu.pl/index.php/PB/article/view/515

50. Gandhi S, Haeussler M, Razy-Krajka F, Christiaen L, Stolfi A. Evaluation and rational design of guide RNAs for efficient CRISPR/Cas9-mediated mutagenesis in Ciona. Developmental Biology. 2017;

51. Kaminow B, Yunusov D, Dobin A, Spring C. STARsolo: accurate, fast and versatile mapping / quantification of single-cell and single-nucleus RNA-seq data. bioRxiv. 2021;1–35.

52. Love MI, Huber W, Anders S. Moderated estimation of fold change and dispersion for RNA-seq data with DESeq2. Genome Biology. 2014;15(12):1–21.

53. Gruber AR, Lorenz R, Bernhart SH, Neuböck R, Hofacker IL. The Vienna RNA Websuite. Nucleic Acids Res [Internet]. 2008 July 1 [cited 2018 July 5];36(Web Server issue):W70–4. Available from: https://www.ncbi.nlm.nih.gov/pmc/articles/PMC2447809/

54. Lewis BP, Burge CB, Bartel DP. Conserved Seed Pairing, Often Flanked by Adenosines, Indicates that Thousands of Human Genes are MicroRNA Targets. Cell [Internet]. 2005 Jan [cited 2018 July 9];120(1):15–20. Available from: http://linkinghub.elsevier.com/retrieve/pii/S0092867404012607

55. Lesage M, Thomas M, Bugeon J, Branthonne A, Gay S, Cardona E, et al. C-ECi: a CUBIC-ECi combined clearing method for three-dimensional follicular content analysis in the fish ovary†. Biology of Reproduction. 2020;103:1099–109.

56. Pachitariu M, Stringer C. Cellpose 2.0: how to train your own model. Nat Methods [Internet]. 2022 Dec [cited 2025 Sept 18];19(12):1634–41. Available from: https://www.nature.com/articles/s41592-022-01663-4

57. Schindelin J, Arganda-Carreras I, Frise E, Kaynig V, Longair M, Pietzsch T, et al. Fiji: an open-source platform for biological-image analysis. Nat Methods [Internet]. 2012 July [cited 2025 Sept 18];9(7):676–82. Available from: https://www.nature.com/articles/nmeth.2019

58. Lesage M, Bugeon J, Thomas M, Pécot T, Thermes V. An end-to-end pipeline based on open source deep learning tools for reliable analysis of complex 3D images of Medaka ovaries. bioRxiv. 2023;2022.08.03.502611.

